# Bacterial LomR Induces the Vibriophage VP882 VqmA-Directed Quorum-Sensing Lysogeny-Lysis Transition

**DOI:** 10.1101/2021.11.15.468771

**Authors:** Jennifer S. Sun, Ameya A. Mashruwala, Chenyi Fei, Bonnie L. Bassler

**Author notes:** Equal contribution (Jennifer S. Sun, Ameya A. Mashruwala).

## Abstract

The bacterial cell-cell communication process called quorum sensing enables groups of bacteria to synchronously alter behavior in response to changes in cell population density. Quorum sensing relies on the production, release, accumulation, and detection of extracellular signal molecules called autoinducers. Here, we investigate a mechanism employed by a vibriophage to surveil host quorum sensing and tune its lysogeny-lysis decision to host cell density. The phage possesses a gene called *vqmA_Phage_* encoding a quorum-sensing receptor homologous to vibrio VqmA. Both VqmA receptors can detect the host bacteria-produced autoinducer called DPO. DPO-bound VqmA_Phage_ launches the phage lysis process. We discover that the bacterial host produces an inducer of the VqmA_Phage_-directed quorum-sensing lysogeny-lysis transition. Production of the inducer appears to be widespread among bacteria. A screen of the *Escherichia coli* Keio collection for mutants impaired for inducer production revealed *lomR*, located in a prophage, and encoding a poorly understood protein. In the *E. coli* screening strain, *lomR* is interrupted by DNA encoding an insertion element. The 3’ domain of this LomR protein is sufficient to induce VqmA_Phage_-directed lysis. Alanine-scanning mutagenesis showed that substitution at either of two key residues abrogates inducer activity. Full-length LomR is similar to the outer membrane porin OmpX in *E. coli* and *Vibrio parahaemolyticus* O3:K6, and OmpT in *Vibrio cholerae* C6706, and indeed, OmpX and OmpT can induce VqmA_Phage_-directed activity. Possibly, development of the LomR, OmpX, or OmpT proteins as tools to direct phage lysis of host cells could be used to control bacteria in medical or industrial settings.

**ABSTRACT IMPORTANCE:** Bacteria communicate with chemical signal molecules using a process called quorum sensing. Quorum sensing allows bacteria to track their cell numbers and orchestrate collective behaviors. Recently, we discovered that a virus that infects and kills bacteria “eavesdrops” on its host’s quorum-sensing process. Specifically, the virus monitors host cell growth by detecting the accumulation of host quorum-sensing signal molecules. In response to the garnered quorum-sensing information, the virus kills the host bacterial cells when the bacterial population has reached a high cell density. This strategy presumably enhances transmission of viruses to new host cells. Here, we discover and characterize three closely-related bacterial host-produced proteins called LomR, OmpX, and OmpT that are capable of inducing the viral quorum-sensing-mediated killing program. Development of this class of inducer proteins as tools to drive “on demand” virus-mediated lysis of pathogenic host bacterial cells could be used to control bacteria in medical or industrial settings.

## INTRODUCTION

The bacterial cell-cell communication process called quorum sensing (QS) enables groups of bacteria to synchronously alter behavior in response to changes in cell population density. QS depends on the production, accumulation, and detection of extracellular signal molecules called autoinducers (AIs) (1, 2). Vibrios are model bacteria used for QS studies. One vibrio QS system is composed of the cytoplasmic AI receptor VqmA and its target, the gene encoding the regulatory small RNA called VqmR (Figure 1). VqmA responds to 3,5-dimethylpyrazin-2-ol (DPO), an AI produced from threonine and alanine. The Tdh (threonine dehydrogenase) enzyme is required for DPO biosynthesis. The DPO-VqmA-VqmR system controls target genes including those required for biofilm formation and virulence (3–5).

**Figure 1.**
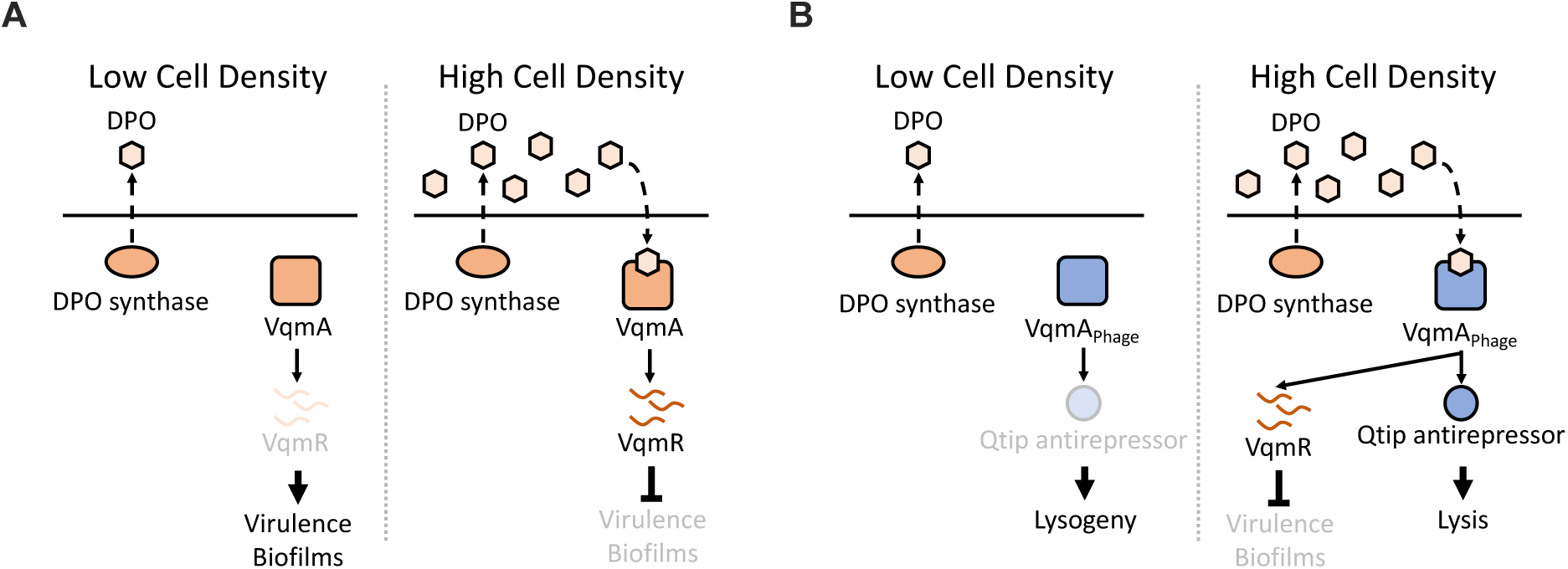
The Phage VP882 Quorum-Sensing (QS) “Eavesdropping” Pathway Controls the Lysogeny-Lysis Fate Decision. Shown is a simplified scheme for vibrio VqmA-directed QS. (**A**, left) At low cell density, the un-liganded VqmA transcription factor has minimal activity, and cells engage in individual behaviors, forming biofilms and producing virulence factors. (**A**, right) At high cell density, VqmA binds to the DPO AI. The complex activates expression of the *vqmR* gene encoding the small RNA VqmR, triggering a cascade that leads to group behaviors, termination of biofilm formation, and repression of virulence factor production. (**B**, left) In phage VP882-infected cells, at low cell density, the un-liganded VqmA_Phage_ transcription factor is inactive, and the phage exists as a lysogen. (**B**, right) At high cell density, VqmA_Phage_ binds to host bacteria- produced DPO. The complex activates expression of the phage *qtip* gene encoding the Qtip antirepressor, which launches the phage lysis program. Also at high cell density, DPO-bound VqmA_Phage_ can activate expression of *vqmR*, presumably altering host QS biology.

A VqmA-type QS system was recently identified in the Myoviridae virus VP882, a non-integrating temperate phage whose natural host is *Vibrio parahaemolyticus*. Phage VP882 infects other vibrios in laboratory settings, including *Vibrio cholerae*, suggesting that a variety of other vibrios could be hosts in nature. The phage VP882 VqmA-type AI receptor is encoded by a gene that we call *vqmA_Phage_*. VqmA_Phage_ “eavesdrops” on vibrio QS (6, 7). Specifically, VqmA_Phage_ binds to host bacteria-produced DPO. Liganded VqmA_Phage_ launches the phage lysis program via production of Qtip, an antirepressor of lysis (Figure 1 and (6, 8)). In addition to inducing *qtip* expression, VqmA_Phage_ can activate host *vqmR* expression (Figure 1), suggesting that the VP882 phage regulates its own lysogeny-lysis decision and host QS behaviors via VqmA_Phage_-directed QS. Other phages also harbor genes encoding putative ligand-binding transcription factors adjacent to genes encoding companion Qtip-type antirepressors (6, 9, 10). These findings suggest that it could be common for phages to control their lysogeny-lysis decisions by tuning into host-produced sensory cues using a Qtip-like lysis de-repression mechanism, and perhaps also alter host biology.

In a previous study, cloned *vqmA_Phage_* that had been reintroduced into the *V. parahaemolyticus* pandemic O3:K6 strain 882, the original host strain from which phage VP882 was isolated, showed that in the absence of DPO, only a basal level of cell lysis occurred, while increased lysis occurred when synthetic DPO was provided (6). This result indicated that DPO drives VqmA_Phage_ activity. Indeed, when *vqmA_Phage_* was eliminated by transposon mutagenesis, DPO-dependent lysis was also eliminated. One caveat of this earlier investigation was that, to facilitate study, *vqmA_Phage_* was artificially induced due to its otherwise low level of expression during lysogeny. Nonetheless, following synthetic induction of phage VP882 *vqmA_Phage_* expression, a positive feedback mechanism increased *vqmA_Phage_* transcription (6). Presumably, this regulatory arrangement enlarges the pool of available VqmA_Phage_ enabling enhanced sensitivity to DPO and, in turn, a more efficient launch of the lysis program. The natural inducer of the VqmA_Phage_-driven lysogeny to lysis switch that initiates the feedback loop was not identified.

Here, we report a class of natural inducers of VqmA_Phage_ activity that are produced by *V. parahaemolyticus*, *V. cholerae,* and other bacteria, including *Escherichia coli*. Using the *E. coli* Keio collection as a screening tool, we identified a mutant defective in production of the inducer, implicating *lomR*. The *E. coli lomR* gene in the Keio collection is interrupted by an IS5Y insertion element (*insH5*) encoding a transposon (11). Our assessment of which region of *lomR’insH5’lomR* is required for inducer activity shows that the DNA encoding the 3’ region of *lomR* (*i.e.*, encoding the C-terminal domain of the LomR protein) is sufficient. Indeed, administration of the purified LomR C-terminal protein fragment to *V. parahaemolyticus* cells induces the phage VP882 VqmA_Phage_-driven lysis program. Alanine scanning mutagenesis of the C-terminal domain of LomR shows that induction activity requires two critical amino acid residues, P8 and D9. LomR resembles outer membrane proteins (OMPs) and is likely of Lambdoid phage origin as *lomR* resides in the *E. coli* Rac prophage (12). OmpX is a LomR homolog present in select Gram-negative bacteria, including in *E. coli* and *V. parahaemolyticus* O3:K6, and the relevant protein is called OmpT in *V. cholerae* C6706. In a test case, we show that *E. coli* OmpX can induce the phage lysis program. Development of LomR or its homologs as tools to drive phage lysis “on demand” could be used to control bacteria in medical or industrial settings.

## RESULTS

### A component present in host cell lysates induces VqmA_Phage_-directed lytic pathway

We investigated whether a bacterial component could be the inducing factor that launches the phage VP882 VqmA_Phage_-directed lysogeny-lysis switch. To test this possibility, we engineered two *E. coli* Top10 reporter strains that harbor *gp69-lux* transcriptional fusions. *gp69* transcription is activated by DPO-bound VqmA_Phage_ and Gp69 is required for phage VP882 VqmA_Phage_-directed host cell lysis. In addition to *gp69- lux*, the *E. coli* reporter strains carried either a phage VP882 with a Tn*5* insertion in an intergenic region (called control phage VP882) or phage VP882 with a Tn*5* insertion in *vqmA_Phage_* (called phage VP882 *vqmA_Phage_*::Tn*5*). The *E. coli* reporter strains naturally make DPO, ensuring DPO is available to bind to VqmA_Phage_. The rationale is to test whether bacteria produce an activity capable of inducing the *gp69-lux* reporter and, if so, whether the component requires VqmA_Phage_ to do so.

In Figure 2A, the first two bars for each reporter strain depict controls and demonstrate how the assay system functions. Administration of LB medium shows the basal levels of *lux* expression that occur in response to endogenously-produced DPO. Administration of LB supplemented with DPO activates expression of the reporter construct, however, only in the strain harboring control phage VP882, demonstrating that endogenously-produced DPO does not saturate the system, and moreover, that DPO- driven induction of *gp69-lux* requires the presence of *vqmA_Phage_*.

**Figure 2.**
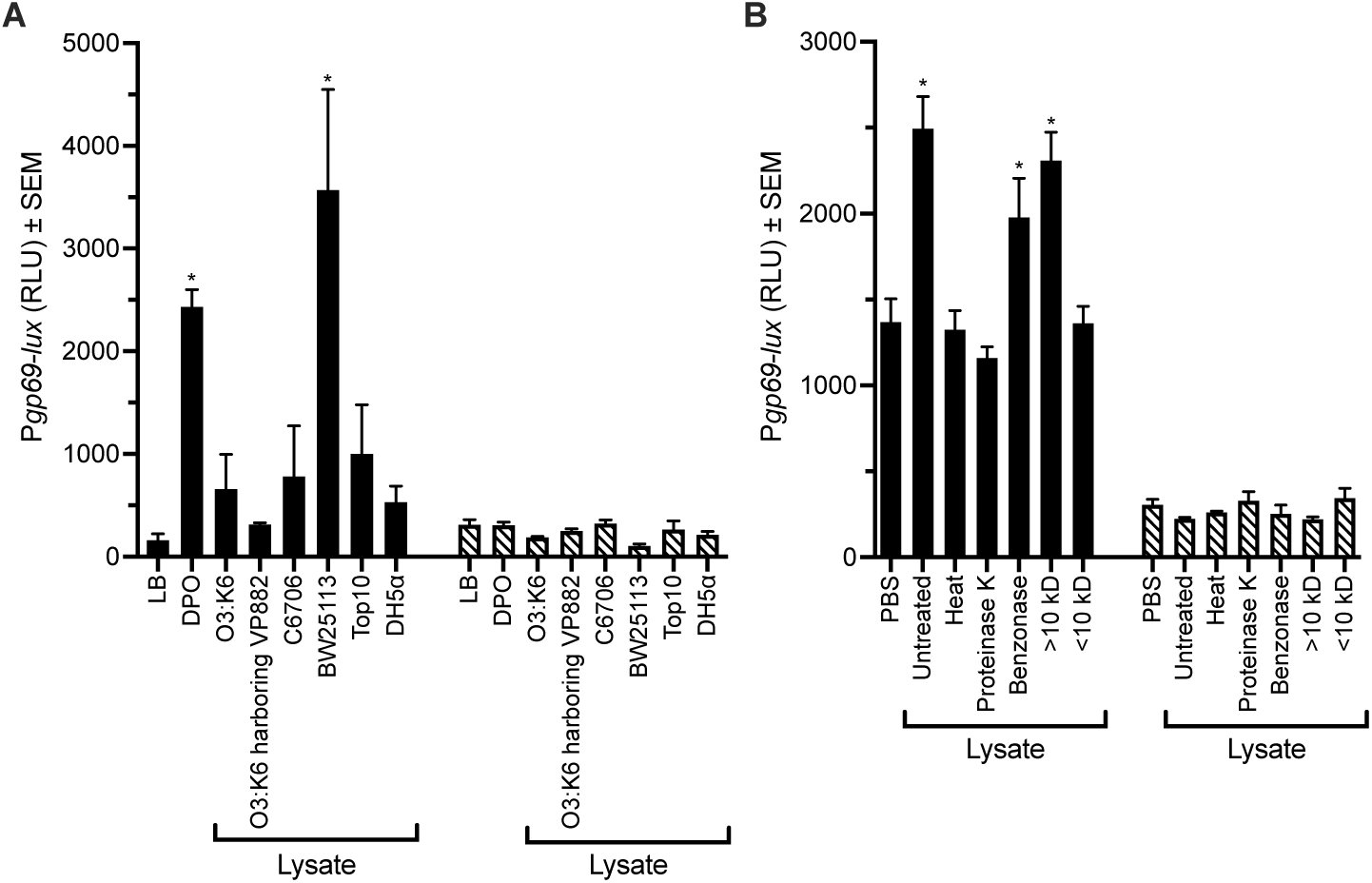
A Bacterial Component Activates the VqmA_Phage_-Directed QS Lysis Cascade. (**A**) Expression of *gp69-lux* in *E. coli* Top10 carrying control phage VP882 (solid bars) or phage VP882 *vqmA_Phage_*::Tn*5* (striped bars) in response to controls – LB medium or 10 μM DPO in LB medium – and to lysates prepared from the designated vibrio (O3:K6 and C6706) and *E. coli* (BW25113, Top10, and DH5*α*) strains. (**B**) Inducing activity in BW25113 lysates treated as follows: untreated, heat; 20 min at 95°C, Proteinase K; 0.8 U for 60 min at 37°C, Benzonase; 1 U for 30 min at 37°C, retentate from 10,000 MWCO filter (labeled > 10 kD), and filtrate from 10,000 MWCO filter (labeled < 10 kD). In this panel, PBS was used as the negative control. Assay and bar patterns as in panel A. Data are represented as mean ± SEM with *n*=4 biological replicates. Statistical significance was calculated using one-way ANOVA Tukey’s multiple comparisons test against (**A**) LB or (**B**) PBS. * denotes *P*<0.05.

To test whether a bacteria-produced component induces VqmA_Phage_-directed activity, and to capture any possible inducer irrespective of type of molecule, we prepared and assessed crude lysates from *V. parahaemolyticus* O3:K6 (hereafter named O3:K6), the natural host of phage VP882. Figure 2A shows that the preparation possesses an activity that modestly induces *gp69-lux* expression only in the *E. coli* reporter strain carrying control phage VP882. Although the data did not show statistical significance, we reproducibly observed low levels of induction from O3:K6 lysates. Likewise, lysate from O3:K6 harboring phage VP882 modestly, but reproducibly, induced *gp69-lux* activity. The fact that lysate from both phage VP882 infected and non-infected O3:K6 displayed some activity reinforces the notion that the inducer is not produced by the phage, but, rather, is a bacterial product. Consistent with this idea, lysates from *V. cholerae* C6706 (hereafter named C6706) as well as strains of *E. coli* from our collection (BW25113, Top10, and DH5*α*) also induced the reporter, albeit to different extents, with the strongest induction by the BW25113 lysate. Together, these data suggest that the inducer is not restricted to vibrios. These data do not allow us to distinguish whether the inducer is identical in the different bacterial genera. We return to this point in a later section. Since lysates from *E. coli* BW25113 exhibit the strongest activation of *gp69-lux* expression among the bacteria we tested we use *E. coli* BW25113 as the source of the inducer activity in most of the remainder of this work. We refer to *E. coli* BW25113 as BW25113 from here forward.

We wondered whether the inducer was a small molecule, DNA, or a protein, as this broad characterization would guide us in its identification. To garner preliminary information, lysate from BW25113 was treated as follows: heat or Proteinase K to denature proteins, Benzonase to degrade DNA, and passage through a 10 kD molecular weight cut-off (MWCO) filter to crudely separate molecules by size (Figure 2B). Heat and Proteinase K treatment eliminated the inducing activity, whereas the activity was resistant to Benzonase. Filtration demonstrated that the factor has a MW > 10 kD. Together, these data indicate that the factor is likely a protein. From here forward we will call this host-produced inducer of the phage VqmA_Phage_ pathway the “inducer protein”.

### Screening of the Keio collection reveals LomR as a candidate protein that induces the phage VP882 QS-directed pathway

BW25113, which makes significant inducer activity (Figure 2A), is the parent strain for an ordered library of *E. coli* null mutants (the Keio collection (13)), providing us a means to identify the inducer protein. Specifically, we screened lysates prepared from the Keio collection strains for those lacking the ability to induce our *E. coli* Top 10 *gp69-lux* reporter. We identified several BW25113 mutants that appeared defective in production of the inducer protein. To verify these results, we reconstructed the mutations in the candidate genes in BW25113. Only lysates from BW25113 lacking *lomR* elicited reduced *gp69-lux* reporter expression compared to lysates from BW25113 (Supplementary Figure 1). Complementation with *lomR* on a plasmid restored inducer production to the BW25113 Δ*lomR* mutant (Figure 3A). Moreover, high level expression of *lomR* resulted in activity in lysates exceeding that naturally made by BW25113 (Supplementary Figure 2). Therefore, BW25113 LomR (named LomR^BW25113^ going forward) appears to harbor the activity that induces the phage VP882 QS-driven lysis pathway.

**Figure 3.**
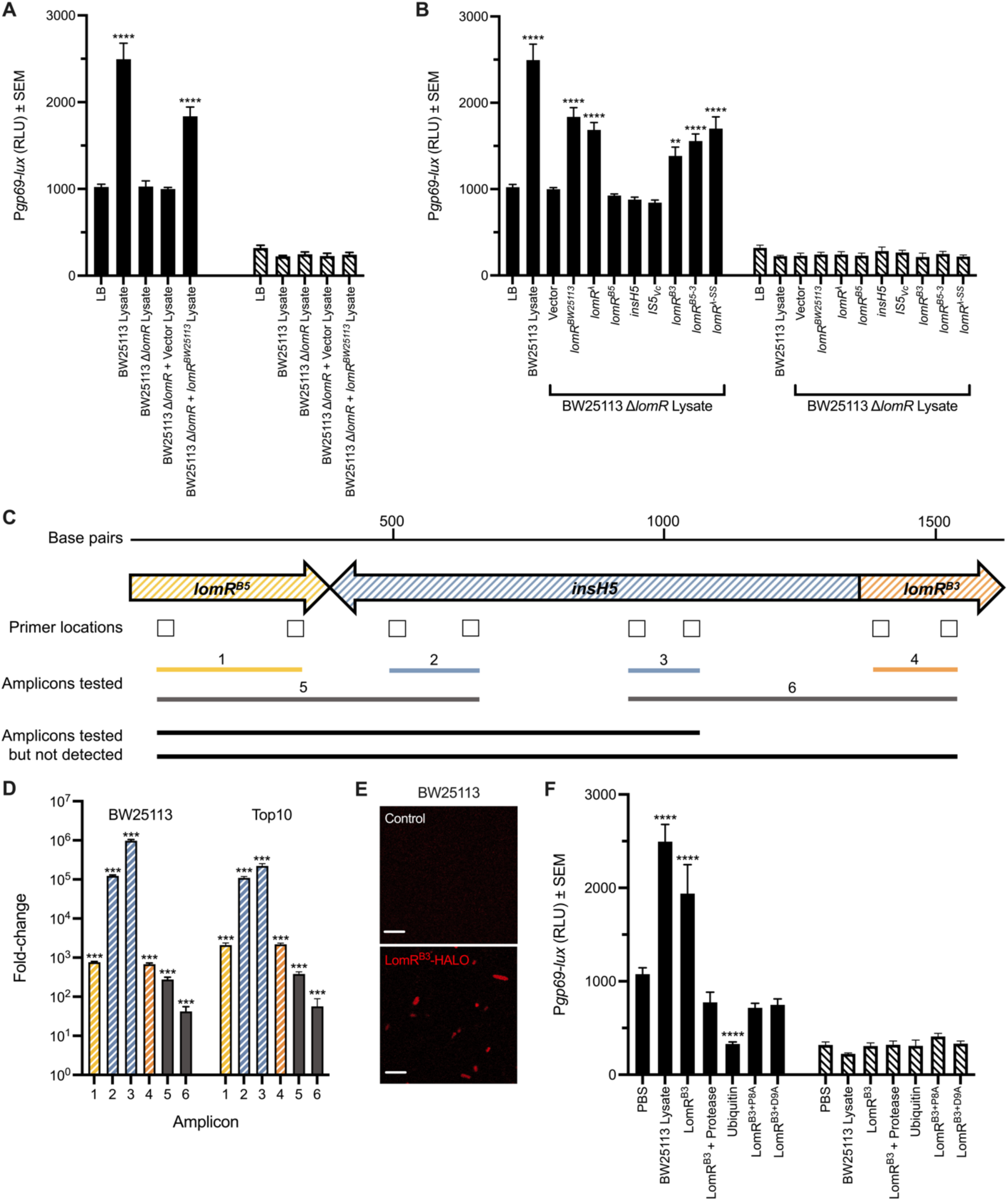
LomR Induces VqmA_Phage_-Directed Activity. (**A**) Inducing activity in lysates prepared from BW25113 Δ*lomR* and BW25113 Δ*lomR* transformed with the pBAD vector (designated Vector) or *lomR* on pBAD (designated *lomR^BW25113^*). (**B**) Lysates prepared from BW25113 and BW25113 Δ*lomR* carrying pBAD (Vector) or the designated genes on pBAD were assayed for activity. (**A,B**) All strains were grown in medium containing 0.2% arabinose prior to lysis. (**C**) Scheme for amplification of regions of *lomR^BW25113^*. The locations of primers (boxes) used for qRT-PCR and the resulting amplicons (numbered lines) are shown. (**D**) Quantitation of qRT-PCR amplification of *lomR^B5^*, *lomR^B3^*, *insH5*, and *lomR^BW25113^* in BW25113 (left) and Top10 (right), represented as fold-change relative to control reactions that lacked reverse transcriptase. Bar colors as in panel C. (**E**) Confocal microscopy of BW25113 lacking LomR^B3^-HALO (upper) and producing LomR^B3^- HALO (lower) visualized with HALO-TMR. Scale bars indicate 6 μm. (**F**) *gp69-lux* expression in *E. coli* Top10 in response to exogenously supplied purified LomR^B3^ or purified LomR^B3^ that had been pre-treated with Proteinase K (designated LomR^B3^ + Protease). Activity following addition of purified LomR^B3^ with alanine substitutions at positions 8 (LomR^B3+P8A^) and 9 (LomR^B3+D9A^). PBS and ubiquitin were added as negative controls and lysate prepared from BW25113 served as the positive control. Data are represented as mean ± SEM with *n*=4 biological replicates. (**A,B,F**) Assay and bar patterns as in Figure 2A. Statistical significance was calculated using one-way ANOVA Tukey’s multiple comparisons test against (**A,B**) LB or (**F**) PBS. Significance in (**D**) was calculated by comparing Ct values from experimental reactions against control reactions lacking reverse transcriptase. ** denotes *P*<0.01 and **** denotes *P*<0.0001.

### A single domain of LomR is sufficient for induction of the phage VP882 QS-directed pathway

*lomR^BW25113^* is encoded in the cryptic Lambda-like Rac prophage (12), which is located downstream of the *trp* operon in many *E. coli* K12 strains (14, 15). *lomR^BW25113^* is most similar to *lomR* in phage Lambda (here designated *lomR^λ^*, Supplementary Figure 3). Phage LomR homologs resemble OMPs belonging to the Gram-negative porin superfamily (11). The functions of phage LomR proteins are poorly understood. What is known is that they are immunogenic porins proposed to assist in small molecule transport, colonization, and adhesion, and they may play roles in cell lysis or in altering permeability of phage infected cells (16, 17). We tested whether *lomR^λ^* could complement BW25113 Δ*lomR* and restore inducer activity. To do this, we transformed BW25113 Δ*lomR* with pBAD*-lomR^λ^*, and assayed lysates made from this strain in our activity assay. There was indeed activity (Figure 3B), suggesting that LomR^λ^ can also function as an inducer.

Often, cryptic prophage genes acquire mutations. Indeed, in BW25113, the *lomR^BW25113^* gene is interrupted by *insH5*, encoding a transposase (InsH) predicted to be involved in transposition of an IS5 insertion element (Supplementary Figure 3 and (23, 29)). The InsH transposase is a member of a family of high-copy number mobile elements that cause topological changes in DNA coupled with transcriptional activation of adjacent genes (19, 20). Because *lomR^BW25113^* contains *insH5*, it cannot simply specify an intact LomR protein. Thus, we wondered what portion(s) of *lomR^BW25113^* is required to generate the inducer activity. The possibilities are: the *lomR^BW25113^* 5’ region upstream of the *insH5* insertion (here designated *lomR^B5^* for <u>B</u>W25113 <u>5</u>’ fragment), *insH5* itself, the 3’ region of *lomR^BW25113^* downstream of the insertion (here designated *lomR^B3^* for <u>B</u>W25113 <u>3</u>’ fragment), an in-frame fusion of the 5’ and 3’ coding regions of *lomR^BW25113^* (here designated *lomR^B5-3^* for <u>B</u>W25113 <u>5</u>’ <u>to</u> <u>3</u>’ fragment), or the entire cassette *lomR*’*insH5*’*lomR* (again, designated *lomR^BW25113^*) (Supplementary Figure 3). We engineered arabinose-inducible versions of each of these regions/genes under control of the pBAD promoter. Additionally, *V. cholerae* possesses a gene called *IS5* (designated here *IS5_Vc_*), a homolog of *insH5*, that is nearly identical in sequence to the *insH5* insertion in *lomR^BW25113^* (76.7% similarity in nucleotide sequence, with 81.9% identity and 90.8% similarity in amino acid sequence) so we also considered the possibility that *IS5_Vc_* could play a role in inducer protein production. To examine this possibility, we cloned *IS5_Vc_* into pBAD. We transformed each of these constructs into BW25113 Δ*lomR*, prepared lysates, and assayed them in our reporter strains. Neither *lomR^B5^*, *insH5*, nor *IS5_Vc_* restored activity. Expression of *lomR^BW25113^, lomR^B3^* and *lomR^B5-3^* complemented the Δ*lomR* defect (Figure 3B). Therefore, only the 3’ domain of LomR is required to restore inducing activity to BW25113 Δ*lomR*. Given that *lomR^BW25113^* contains an apparently inactivating insertion, we wondered how we identified *lomR^BW25113^* in our Keio collection screen. Analysis of the DNA sequence in this region shows that in *lomR^BW25113^*, downstream of the *insH5* insertion, is DNA encoding a start codon that lies in frame with the DNA encoding LomR^B3^ (Supplementary Figure 3). Presumably, this start codon is employed to produce LomR^B3^ in BW25113. Indeed, qRT-PCR confirmed that *lomR^B5^*, *insH5*, and *lomR^B3^* are expressed in BW25113 (Figure 3C-D) with *insH5* being the most highly expressed and *lomR^B5^* and *lomR^B3^* expressed at similar levels. We could not amplify full-length *lomR^BW25113^* region suggesting no transcription of the full region occurs due to disruption by *insH5* (Figure 3C- D). We obtained similar results in *E. coli* Top10 (Figure 3C-D), the strain we used in our reporter assays that also exhibits inducing activity (Figure 2A) and has an *insH5* element inserted in *lomR*.

As above, we carried out qRT-PCR with the BW25113 Δ*lomR* strain. Unexpectedly, the strain produced *lomR^B5^* and *insH5* transcripts, but it did not produce detectable *lomR^B3^* transcript (Supplementary Figure 4). The BW25113 Δ*lomR* strain used in our study was reconstructed from the original Keio mutant by P1 phage transduction of the *lomR*::*kan* region into BW25113 followed by elimination of the kanamycin resistance marker (see Methods). Examination of the primer sequences used by Baba *et. al* (13) to construct the Δ*lomR* strain in the Keio collection revealed that their design deleted the 3’ portion of the *lomR^BW25113^* locus leaving the upstream sequence (*i.e.,* the region that we call *lomR^B5^*) and a majority of the *insH5* sequence intact. Thus, we conclude that the original Keio mutant and the BW25113 Δ*lomR* strain we made and used in our study lacks only *lomR^B3^*. These results are consistent with our finding that, in this locus, expression of only the *lomR^B3^* gene segment is required for inducer activity.

In *E. coli* K12 strains that encode intact LomR proteins and/or that have LomR^λ^, these proteins possess N-terminal signal sequences that presumably drive localization to the outer membrane, consistent with their being OMPs. However, LomR^B3^ in BW25113 does not have a signal sequence and lysates prepared from strains in which we introduce DNA encoding only LomR^B3^ exhibit activity (Figure 3B). Thus, localization of LomR^B3^ to the outer membrane appears dispensable for activity. To verify this notion, we focused on LomR^λ^, which we have shown can substitute for LomR^B3^ (Figure 3B). We engineered arabinose-inducible *lomR^λ^* with its native signal sequence removed (called *lomR^λ-SS^*). Lysates prepared from BW25113 Δ*lomR* carrying *lomR^λ-SS^* activated the reporter (Figure 3B). With respect to LomR^B3^, we fused it to Halo, introduced it into BW25113, and showed by confocal microscopy that LomR^B3^-Halo is indeed diffuse in the cell (Figure 3E). Together, these findings suggest that LomR^B3^ inducer activity does not depend on transport to and insertion into the outer membrane. All of these results are consistent with our crude lysate preparations containing both membrane and cytoplasmic contents (see Methods).

To garner additional proof that activation of the VqmA_Phage_-directed lysogeny to lysis transition can be induced by LomR^B3^, we purified this domain and added it exogenously to the *E. coli* Top10 *gp69-lux* reporter strains. Purified LomR^B3^ was sufficient for induction of activity (Figure 3F). Consistent with this finding, pre-treating purified LomR^B3^ with Proteinase K eliminated inducer activity (Figure 3F). As a control, we also supplied ubiquitin, a protein similar in size to LomR^B3^, to the reporter strain. Ubiquitin did not induce *gp69-lux* (Figure 3F). To pinpoint the amino acid (AA) residues in LomR^B3^ required to confer inducer activity, we used alanine scanning mutagenesis. LomR^B3^ is 86 AAs in length. Initially, we mutated every set of three contiguous AAs to alanines, revealing the A7-P8-D9 stretch as implicated in inducer activity (Supplementary Figure 5). Indeed, individual substitutions at either residue 8 or 9 abrogated the inducer activity of the purified proteins (Figure 3F). The residue at position 7 is naturally an alanine.

### LomR and its homologs are host-produced proteins capable of inducing VqmA_Phage_- directed lysis

LomR homologs exist in prophages and cryptic phages in Gram-negative bacteria beyond *E. coli* K12 strains (Figure 4A-C). Among similar proteins that are not of phage origin, sequence alignment shows that OmpX is the closest homolog (Figure 4B). We wondered if OmpX possesses inducer activity, and if so, whether activity is a general feature of OMPs. BW25113 encodes 10 OMPs – OmpA, OmpC, OmpE, OmpG, OmpH, OmpL, OmpN, OmpT, OmpW, and OmpX. We individually deleted the genes encoding each OMP. Figure 4D shows that lysate from only the BW25113 strain lacking OmpX lost the capability to activate *gp69-lux*. Complementation with cloned *ompX^BW25113^* rescued inducer activity (Figure 4E). These results suggest that the *vqmA_Phage_* pathway can be activated by LomR^B3^ and/or its homolog OmpX^BW25113^, but not generally by OMP proteins. We do not understand why *ompX^BW25113^* was not revealed in our original screen of the Keio collection.

**Figure 4.**
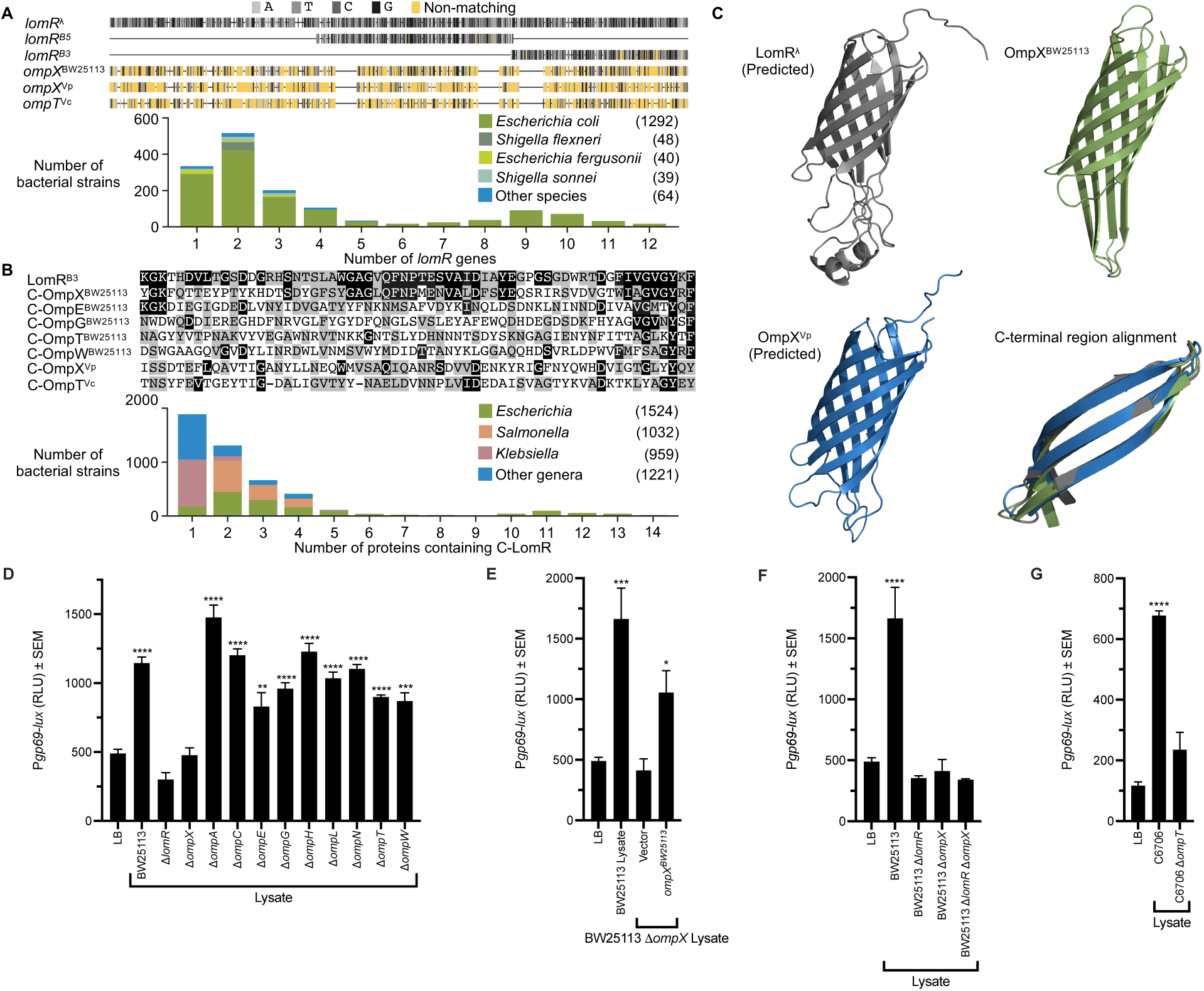
LomR Conservation and OmpX and OmpT as Inducers of the VqmA_Phage_- Directed Lysogeny-Lysis Transition. (**A**, top) Multiple sequence alignment of the indicated genes. Grayscale and yellow shading indicate identical and non-identical nucleotides, respectively. All alignments are relative to *lomR*^λ^. Horizontal bars indicate gaps. (**A**, bottom) Histogram showing the number of *lomR* genes per bacterial strain predicted by DNA sequence homology. Colors designate bacterial species. Values in parentheses indicate the number of bacterial strains that possess at least one *lomR* gene for the designated species. (**B**, top) Multiple sequence alignment of the C-terminal regions of the indicated proteins. Black and gray indicate, respectively, identical and similar amino acids according to the substitution matrix BLOSUM62. Residues P8 and P9 that we find are critical for *E. coli* LomR^B3^ function (see Supplementary Figure 5) are not conserved in all LomR proteins. (**B**, bottom) Histogram showing the number of proteins containing the C-terminal region of LomR per bacterial strain predicted by protein sequence homology. Colors indicate bacterial genera. Values in parentheses indicate the number of bacterial strains that possess at least one protein containing the C-terminal region of LomR for the designated genera. (**A,B**) Vp and Vc denote genes or proteins in *V. parahaemolyticus* and *V. cholerae,* respectively. (**C**) Predicted and actual structures of (i) LomR^λ^, (ii) OmpX^BW25113^, and (iii) OmpX^Vp^; and (iv) the structural alignment of their C- terminal 46 amino acids. (i) and (iii) are predicted structures by HHpred (see Materials and Methods). (**D**) Activities in lysates prepared from BW25113 and BW25113 harboring deletions in the designated genes. (**E**) Activity in lysates prepared from BW25113 and BW25113 Δ*ompX* complemented with arabinose-inducible *ompX^BW25113^*. (**F**) Activity in lysates prepared from BW25113, BW25113 Δ*lomR*, BW25113 Δ*ompX*, and BW25113 Δ*lomR* Δ*ompX*. (**G**) Activity in lysates prepared from C6706 and C6706 Δ*ompT*. (**D-G**) Data are represented as mean ± SEM with *n*=4 biological replicates. Assay in D-G as in Figure 2A. Statistical significance was calculated using one-way ANOVA Tukey’s multiple comparisons test against LB. * denotes *P*<0.05, ** denotes *P*<0.01, *** denotes *P*<0.001, and **** denotes *P*<0.0001.

Not surprisingly, lysate from the BW25113 double Δ*lomR* Δ*ompX* mutant possessed no inducer activity, consistent with the inability of lysates prepared from the single Δ*lomR* and single Δ*ompX* mutant strains to induce the reporter (Figure 4F). Together, the results with the three mutants suggest that either OmpX^BW25113^ or LomR^B3^ is sufficient for inducer production. Because their activities are not additive, an epistatic relationship could exist between LomR^B3^ and OmpX^BW25113^ in which one of the proteins controls the other’s production or activity. To examine this possibility, we used qRT-PCR to measure transcription of *lomR^B^*^3^ and *ompX^BW25113^* in both single mutants. Supplementary Figure 6 shows that deletion of *lomR^B3^* or deletion of *ompX^BW25113^* did not affect transcription of, respectively, *ompX^BW25113^* or *lomR^B3^*. Thus, we do not find evidence for an epistatic connection. Our data do not eliminate the possibility that the LomR^B3^ and OmpX^BW25113^ proteins may interact with and alter one another’s activity.

*V. parahaemolyticus* and *V. cholerae* do not possess *lomR* genes, however, they do harbor *ompX*-type genes. The C6706 *ompX* homolog is called o*mpT* (Figure 4A-C). Thus, OmpX in O3:K6 (OmpX^Vp^) and OmpT in C6706 (OmpT^Vc^) could act as inducers of the phage VP882 lysis cascade. To test this possibility, we deleted *ompT^Vc^* from C6706. O3:K6 is not amenable to genetic manipulation so we could not test function in this strain. Figure 4G shows that lysate prepared from C6706 Δ*ompT* does not exhibit inducer activity in contrast to lysate prepared from WT C6706, confirming that OmpT^Vc^ can also act as an inducer of VqmA_Phage_-directed activity.

### LomR and OmpX can induce lysis in phage VP882-infected V. parahaemolyticus, the natural host for phage VP882

Our above studies relied on a heterologous reporter as a proxy for VqmA_Phage_- directed lysis. To test whether purified LomR^B3^ and purified OmpX^BW25113^ are capable of inducing lysis *in vivo*, we assessed *gp69-lux* expression, and simultaneously, we measured cell lysis by tracking optical density of O3:K6 carrying control phage VP882. As a control, we administered mitomycin C (MMC), a known inducer of lysis (6, 7). MMC activated *gp69-lux* and drove cell lysis (Figure 5A and 5B, respectively). Exogenous addition of purified LomR^B3^ or purified OmpX^BW25113^ to O3:K6 caused equally strong activation of the *gp69-lux* fusion (Figure 5A) and drove moderate cell lysis when phage VP882 was present (Figure 5B). Protease pretreatment of LomR^B3^ and OmpX^BW25113^ eliminated lysis capability. In the opposite vein, addition of ubiquitin did not induce reporter expression or lysis (Figure 5A and 5B, respectively).

**Figure 5.**
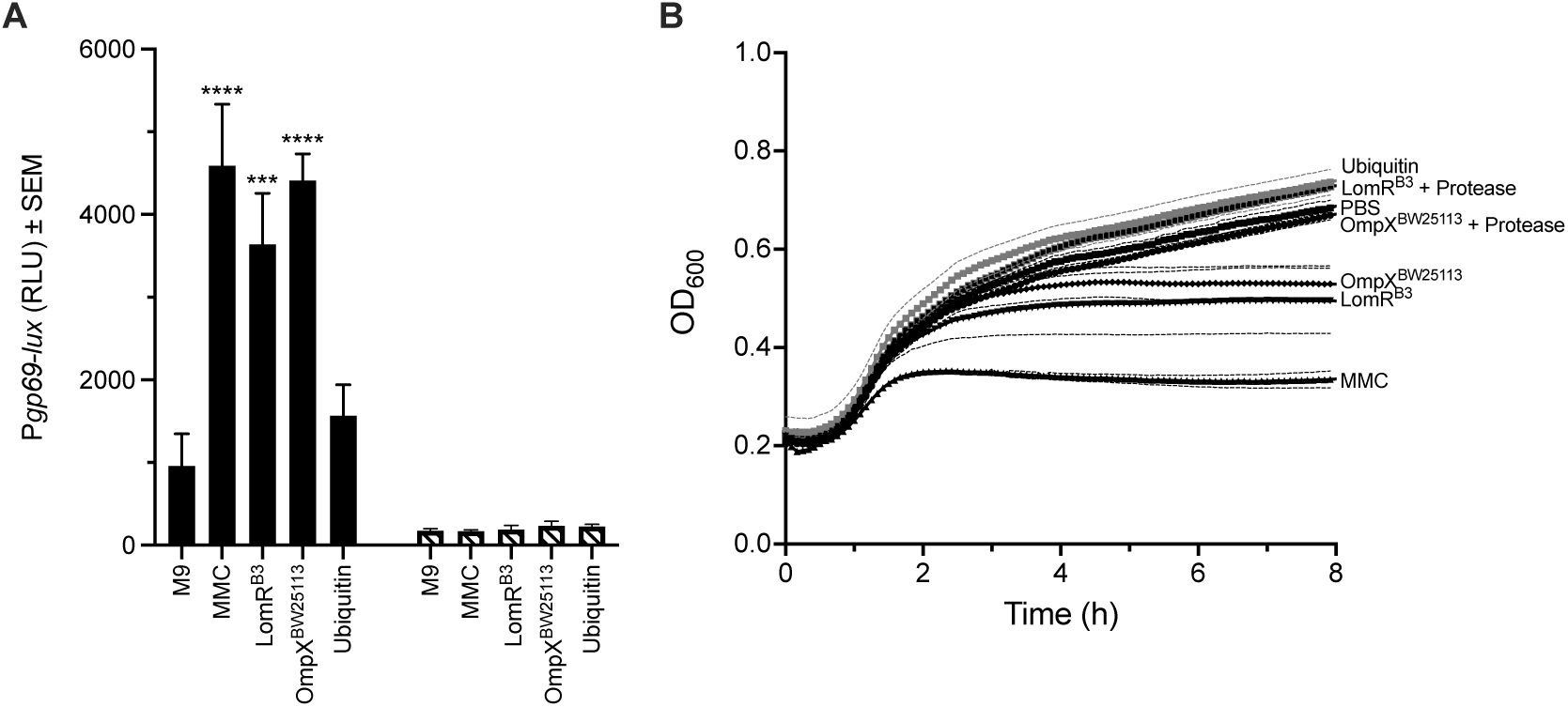
LomR and OmpX Can Induce Lysis in Phage-Infected *V. parahaemolyticus* O3:K6. (**A**) Activity of the *gp69-lux* reporter in O3:K6 in response to the designated treatments. M9 is the M9 medium control. MMC is mitomycin C. Purified LomR^B3^, OmpX^BW25113^, or Ubiquitin were added at 10 μM. Assay and bar patterns as in Figure 2A. (**B**) Growth of O3:K6 harboring phage VP882 treated with the designated purified proteins or with the designated purified proteins that had been pre-treated with protease. PBS buffer was used as the control. Data are represented as mean ± SEM (dashed lines in B) with *n*=4 biological replicates. In (**A**), statistical significance was calculated using one-way ANOVA Tukey’s multiple comparisons test against M9. *** denotes *P*<0.001, and **** denotes *P*<0.0001.

All of the preceding assays relied on exogenous addition of the LomR^B3^ (or OmpX^BW25113^) inducer to cells strongly suggesting that LomR^B3^ acts from the extracellular environment. As a preliminary test of whether the VqmA_Phage_-directed lysis pathway can be induced by LomR^B3^ that is present in the cytoplasm, we overexpressed *lomR^B3^* on pBAD in BW25113 Δ*lomR* carrying either control phage VP882 or phage VP882 *vqmA_Phage_*::Tn*5* and examined whether lysis occurred. Supplementary Figure 7 shows no difference in cell growth following *lomR^B3^* overexpression. Consistent with this finding, we note that *E. coli* Top10, the strain we use for our reporter assay, expresses *lomR^B3^* at a level similar to that expressed by BW25113 (Figure 3D), yet, cell lysis does not occur. This observation further suggests that physiological levels of LomR^B3^ in the cytoplasm are insufficient for activation of the VqmA_Phage_ pathway and LomR^B3^ must be provided from the external environment. Thus, to date, we have no evidence that LomR can function from the cytoplasm.

## DISCUSSION

The DPO-VqmA QS AI-receptor coordinates group behaviors in vibrios (Figure 1). A homologous VqmA, called VqmA_Phage_, encoded in vibriophage VP882 enables the phage to eavesdrop on and manipulate its vibrio host’s QS-mediated communication process. The cue that naturally activates the VqmA_Phage_-directed lysis cascade was unknown. Here, we discovered that the bacterial LomR protein and its homologs OmpX and OmpT can function as inducers (Figures 3-5).

LomR homologs are encoded in cryptic phages in bacterial genomes, suggesting that LomR has a phage origin (Figure 4). Consistent with this notion, phage Lambda LomR, which shares sequence similarity with *E. coli* BW25113 LomR, can complement an *E. coli* Δ*lomR* strain to induce VqmA_Phage_-directed activity. Curiously, only the C-terminal domain of BW25113 LomR is required to induce VqmA_Phage_ activity. Precedent for functional C-terminal domains of OMPs exists. For example, the C-terminus of *E. coli* OmpC is superior to full-length OmpC at activating the DegS protease to launch the envelope stress response. The difference in activity stems from stronger binding of truncated OmpC than full-length OmpC to DegS (18). Nonetheless, full-length LomR and OmpX proteins localize to the bacterial outer membrane (18). However, it cannot be that translocation to the membrane is required for induction of the VqmA_Phage_-directed lysis pathway. We say this because, at least in BW25113, the LomR protein is naturally truncated and possesses no signal sequence. Likewise, lysates prepared from cells producing Lambda LomR lacking its signal sequence induced the VqmA_Phage_-directed lysis pathway (Figure 3B). These observations are consistent with our finding that LomR (truncated and full-length) and OmpX induce VqmA_Phage_ activity from the extracellular environment (Figures 2, 3, and 5). Possibly, inducer protein residing in the outer membrane (in the case of full-length proteins) or residing in the cytoplasm (in the case of truncated versions and prior to transport to the outer membrane) are released as a consequence of ordinary (*i.e.*, non-phage-mediated) cell lysis and provide the source of extracellular inducer in nature. We do not yet understand the mechanism underlying LomR^B3^/OmpX^Vp^/OmpT^Vc^ activation of the phage lysis program. Possibilities include increased *vqmA_Phage_* expression, increased VqmA_Phage_ production, increased VqmA_Phage_ activity, or increased DPO production. We are currently exploring the underlying mechanism.

The phage Lambda LomR porin protein has long been considered an accessory protein because it is not essential for phage replication under laboratory conditions (21). Rather, Lambda LomR is thought to function in lysogenic conversion. LomR proteins that are encoded on cryptic phages residing in bacterial genomes are proposed to provide adaptative advantages to the host bacteria, particularly during pathogenesis (21, 22). For example, in *Brucella* species, LomR increases mammalian host cell adhesion, invasion, and evasion of mammalian host defenses (23). In some disease-causing *E. coli* strains, the LomR protein (in these cases full-length LomR, not truncated LomR) produced from the genome is implicated as a virulence factor that triggers bacterial attachment to human buccal epithelial cells, a step required for infection (21). Regarding origins, phage Lambda LomR is presumed to be the ancestor of a five-member family of virulence-related outer membrane proteins in Gram-negative bacteria, including Ail (24), Omp21 (25), PagC (26), Rck (27), and OmpX (28). All of these proteins share highly conserved residues in their *β*-barrel cores (29). Our results suggest that perhaps phage VP882 exploits this family of Lambda-derived LomR homologs to synchronize its lysogeny to lysis transitions with high host cell density. Such a strategy could improve phage VP882 dissemination.

We suggest a model in which a class of bacterial host factors regulate this phage pathway, which is surprising given that induction drives host cell killing by the phage. *V. parahaemolyticus* containing phage VP882 exists in nature, and in our laboratory analyses, the cells do not die at high cell density. Thus, a mechanism must operate to restrict VqmA_Phage_ activity and enable lysogeny. Perhaps, in *V. parahaemolyticus*, the requirement for OmpX-mediated induction of VqmA_Phage_ activity provides this mechanism. In the case of lysogens, mechanisms need to exist to tamp down OmpX production and/or OmpX export to enable later VqmA_Phage_ function when high host cell density is achieved and lysis is warranted. How these different phage lifestyles are maintained, how transitions between them occur, and the precise role(s) OmpX plays in phage lifestyle decision making events await future analyses.

Known inducers of phage lysis consist almost exclusively of synthetic (*i.e.,* laboratory-used) DNA-damaging agents such as MMC that do not exist in authentic environments in which bacteria and phages co-reside. Our discoveries suggest that in *V. parahaemolyticus*, OmpX could function as a natural regulator that dictates the timing of the phage lysis program. We speculate that under particular conditions, OmpX accumulates in the extracellular environment as cell density increases, perhaps in step with accumulation of the DPO AI, and together or sequentially, DPO and OmpX induce the phage to switch from lysogeny to lysis. More certain is that the QS regulatory arm of this phage’s lysogeny-lysis transition can be governed by two bacteria-derived products, DPO and OmpX, suggesting that discovering how non-traditional stimuli affect lysogeny-lysis pathways could improve our general understanding of how phages make decisions in real-world environments. Further studies are required to determine if LomR and its homologs, such as OmpX and OmpT, can induce lysis in natural systems. If so, one can imagine them being developed as tools to drive phage lysis “on demand,” to control pathogenic bacteria in medical or industrial contexts.

## MATERIALS AND METHODS

### Bacterial Strains

Strains used in this study are listed in Table S1A. All strains were grown with shaking at 250 rpm. Vibrio strains were grown at 30°C in Luria Marine (LM) medium or in M9 medium supplemented with 0.5% glucose and 0.4 mM amino acids. *E. coli* strains were grown at 37°C in Luria Broth (LB). Antibiotics were used at the following concentrations (μg mL^-1^): ampicillin, 200; chloramphenicol, 10; kanamycin, 100. Induction of genes cloned onto pBAD was achieved by the addition of 0.2% arabinose (Sigma), unless otherwise indicated.

### Strain Construction

Primers and gene blocks were obtained from Integrated DNA Technologies and are listed in Tables S1B and S1C, respectively. Plasmids used in this study were validated by sequencing (Genewiz) and are listed in Table S1D. PCR with Phusion High Fidelity DNA Polymerase (NEB) or iProof DNA Polymerase (Bio-Rad) was used to generate inserts and backbone DNA for Gibson assembly (NEB, NEBuilder HiFi DNA Assembly mix), yeast recombination-assisted assembly (30), and traditional cloning (NEB, restriction enzymes and T4 DNA ligase). Constructs were initially transformed into *Saccharomyces cerevisiae* FY2 (30) or *E. coli* Top10 (Invitrogen). DNA was introduced by electroporation using 0.1 cm gap cuvettes (USA Scientific) with a Bio-Rad MicroPulser. In-frame deletions of candidate genes encoding the inducer were generated via phage P1 transduction (31) followed by Flp-catalyzed excision of the kanamycin cassette (32).

### Lysate Preparation

Bacterial strains were inoculated from single colonies and grown overnight at 37°C with shaking at 250 rpm. These cultures were back-diluted 1:100 with LB to 50 mL, followed by growth at 37°C with shaking at 250 rpm to OD_600_ = 1.0. The cultures were placed on ice after which the samples were divided into aliquots of 10 mL, placed in 15 mL conical tubes, and the cells immediately pelleted at 4,000 rpm for 10 min at 4°C. Pellets were rinsed twice with 10 mL distilled water. The cells were then resuspended in 10 mL 1x phosphate-buffered saline (PBS) pH 7.4 containing protease inhibitor cocktail (MilliporeSigma, cOmplete Mini EDTA-free) and subjected to three freeze-thaw cycles with liquid nitrogen followed by sonication. A brief centrifugation step (1,000 rpm for 5 min at 4°C) removed large debris. These lysates preparations were always used within 5 h.

### Bioluminescence Assays

Assays were performed as described in Silpe & Bassler 2019 (6) with the following modifications. 250 ng mL^-1^ MMC, 10 μM DPO, 10 μL lysate, or 10 μM purified protein was added as specified. Wells that did not receive treatments were provided equivalent volumes of vehicle corresponding to the treatment (LB for lysates, M9 for DPO, and PBS for MMC and purified proteins). Single time point assays were monitored at 3.5 h.

### Protein Purification

The genes encoding 6xHIS-LomR^B3^, 6xHIS-LomR^B3+P8A^, 6xHIS-LomR^B3+D9A^, or 6xHIS- OmpX^BW25113^ were cloned into pET28b and overexpressed in *E. coli* BL21(DE3). Cultures were inoculated from single colonies and grown overnight at 37°C with shaking at 250 rpm. These cultures were back-diluted 1:100 with LB to 500 mL, followed by growth at 37°C with shaking at 220 rpm to OD_600_ = ∼0.5-0.8. The temperature was reduced to 18°C for 1 h prior to addition of 1 mM isopropyl *β*-D-1-thiogalactopyranoside (IPTG) and incubation overnight. The cultures were placed on ice and triplicate samples were pelleted at 4,000 rpm for 10 min at 4°C. Cell lysis was accomplished by resuspending each pellet in 20 mL of lysis buffer (25 mM Tris-HCl pH 8.0, 150 mM NaCl, 0.1 mM protease inhibitor cocktail, and 0.5 mM DTT), followed by a freeze-thaw cycle in liquid nitrogen, sonication on ice, and centrifugation at 13,000 rpm for 1 h at 4°C. 2 mL of nickel-charged resin (Qiagen, Ni-NTA Agarose) was added to a column (Qiagen, 5 mL polypropylene column), equilibrated with lysis buffer, and maintained at 4°C. The clarified supernatants from the lysed cells were loaded onto the nickel resin, washed with 40 mL wash buffer (25 mM Tris-HCl pH 8.0, 300 mM NaCl, 20 mM imidazole, and 0.5 mM DTT), and eluted with 20 mL of elution buffer (25 mM Tris-HCl pH 8.0, 150 mM NaCl, 300 mM imidazole, and 0.5 mM DTT). The eluate was concentrated to 1 mL using 10 kD MWCO centrifuge filters (MilliporeSigma, Amicon Ultra-15), followed by separation on a Superdex-200 size exclusion column (GE Healthcare) in gel filtration buffer (25 mM Tris-HCl pH 8.0, 100 mM NaCl, and 2 mM DTT). Protein was concentrated, flash frozen, and stored at -80°C.

### qRT-PCR

Overnight cultures of BW25113 and Top10 were diluted 1:1000 and grown to OD_600_ = 2.0 prior to harvest. Transcripts were stabilized and preserved with RNA-Protect reagent (Qiagen) as recommended by the manufacturer. RNA was extracted and processed for qRT-PCR as described (33). In the cases in which no transcripts were detected in control reactions lacking reverse transcriptase, for calculation purposes, the Ct value was artificially set to 40, which is the maximum number of cycles used in the PCR reaction and thus represents the limit of detection for our sample set.

### Confocal Microscopy

Live-cell staining of BW25113 carrying HALO fusions was performed using a previously described HALO-TMR protocol described (6) with the following modification: 1-5 μL of samples were spotted onto glass slides, sandwiched between glass coverslips, and excess moisture wicked away to fix the cells.

### Bioinformatic Analyses

#### Sequence homologs search

To identify regions of similarity between DNA or protein sequences, local sequence alignments were performed in MATLAB (Mathworks, 2020) using the Smith-Waterman (SW) algorithm (34).

##### Query sequence

To identify bacterial genes encoding LomR family proteins, *lomR^B3^* was used as the query. The *lomR^B3^* sequence is highly homologous to the 3’-most 190 nucleotides of *lomR^λ^* (Figure 4A). The amino acid sequence encoded by *lomR^B3^* was used as the query for protein sequence homology analyses.

##### Search set

21,076 bacterial strains with fully assembled genome sequences were scanned. Their genomic DNA sequences were downloaded from the GenBank database (35).

##### Similarity scoring

For DNA sequence alignments using the SW algorithm, identical nucleotides were assigned a score of +1, and non-identical nucleotides or gaps were assigned a score of -2. For protein sequence alignments, the standard substitution matrix BLOSUM62 was used to compute similarity scores (36). The *p*-values associated with the similarity scores that the SW algorithm yielded were computed by applying a shuffling algorithm to the subject sequences. A *p*-value of < 10^-6^ was used to identify homologous sequences. Sequence homologs were annotated by PROKKA (37) and were verified to encode LomR family or OmpX family proteins.

#### Protein structural analyses

HHpred (MPI Bioinformatics Toolkit) was used to identify bacterial proteins possessing structural similarity to the query proteins (38). The top ∼8 to 10 hits from the search were used as templates to predict the structure of the query protein using MODELLER software (39). Visualization and spatial alignments of protein structures were performed using PyMOL (Schrödinger, version 2.4.2).

### Quantification and Statistical Analysis

Data are presented as the mean ± standard errors of the means. The number of independent biological replicates for each experiment is indicated in the figure legends. No blinding or randomization was used in these experiments. All bioluminescence data were recorded, analyzed, and plotted using Microsoft Excel and GraphPad Prism 9. Imaging data were collected, processed, and prepared using LASX and Fiji.

### Data Availability

All experimental data that support the findings of this study are available from the corresponding author upon request.

## ACKNOWLEDGEMENTS

We thank members of the Bassler laboratory for insightful discussions. This work was supported by the Howard Hughes Medical Institute, National Institutes of Health Grant R37GM065859, and National Science Foundation Grant MCB-1713731 (BLB), National Institutes of Health Grant F32GM139233 (JSS), and Life Sciences Research Foundation Award sponsored by the Howard Hughes Medical Institute (AAM). The content is solely the responsibility of the authors and does not necessarily represent the official views of the National Institutes of Health.

---

**Supplementary Figure 1.**
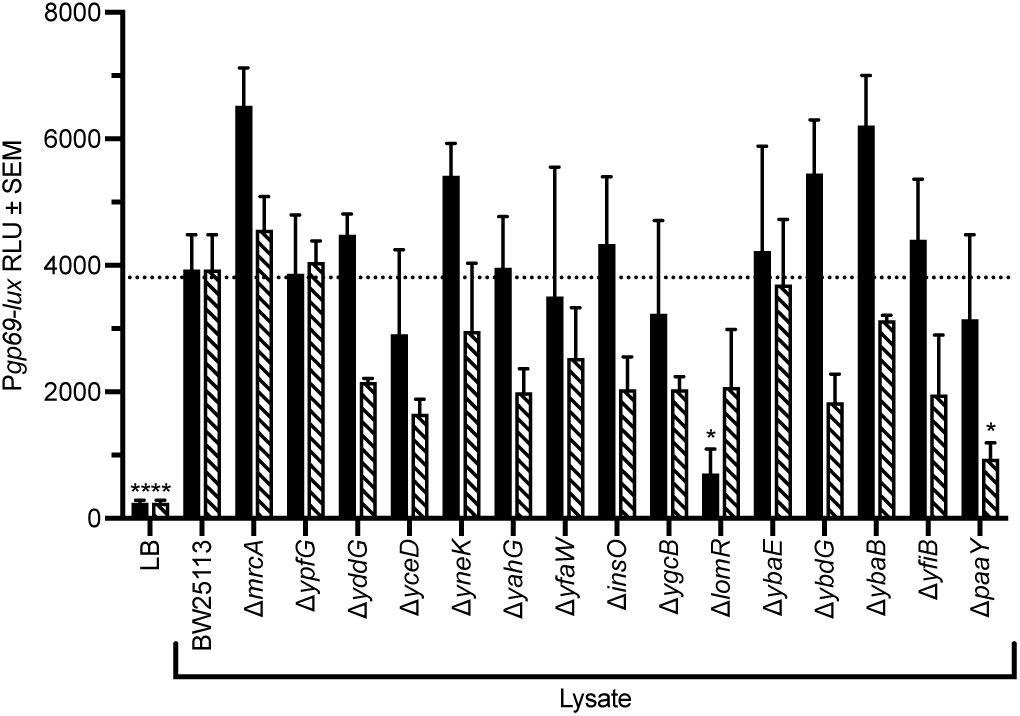
Keio Collection Screen for Mutants Defective in Production of the Protein that Induces the VqmA_Phage_-Directed Pathway. Activity from lysates prepared from BW25113 and deletion strains derived from candidate Keio collection mutants with putative defects in production of the inducer protein. Assay and bar patterns as in Figure 2A. Data are represented as mean ± SEM with *n*=8 biological replicates. The dotted line denotes the average level of activity from the BW25113 lysate. Striped bars are plotted for the LB and *E. coli* BW25113 control experiments, but because the values are low, they are difficult to see on the graph. Statistical significance was calculated using one-way ANOVA Tukey’s multiple comparisons test against BW25113 lysate. * denotes *P*<0.05, and ** denotes *P*<0.01.

**Supplementary Figure 2.**
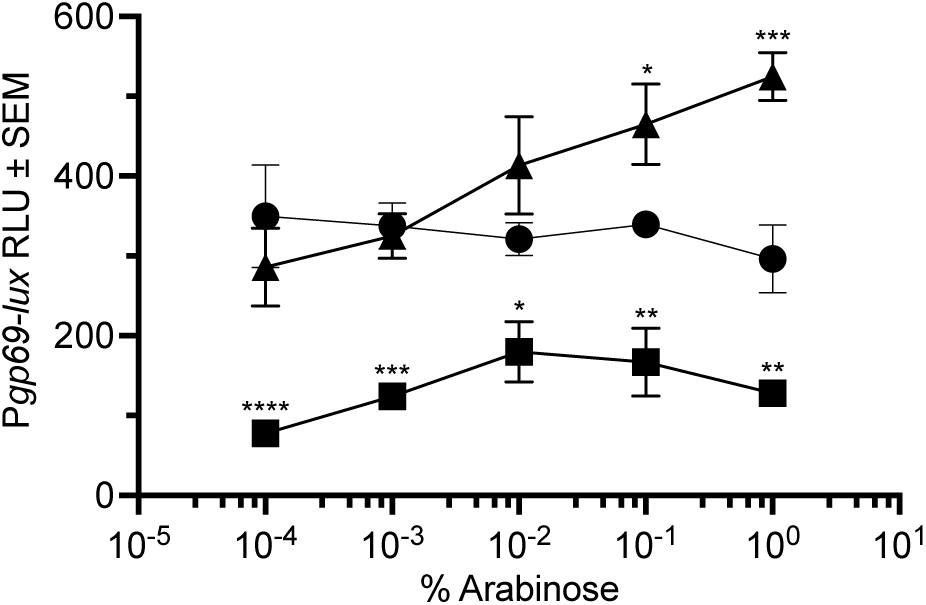
Overexpression of *lomR^BW25113^* Increases Production of Inducer Activity. Expression of *gp69-lux* in *E. coli* Top10 carrying control phage VP882 in response to lysates from BW25113 (circles), BW25113 Δ*lomR* (squares), or BW25113 Δ*lomR* carrying cloned *lomR^BW25113^* (triangles). Strains were grown for 4 h in the presence of the specified concentrations of arabinose prior to lysis. Data are represented as mean ± SEM with *n*=4 biological replicates. Statistical significance was calculated using one-way ANOVA Tukey’s multiple comparisons test against BW25113 lysate. * denotes *P*<0.05, ** denotes *P*<0.01, *** denotes *P*<0.001, and **** denotes *P*<0.0001.

**Supplementary Figure 3.**
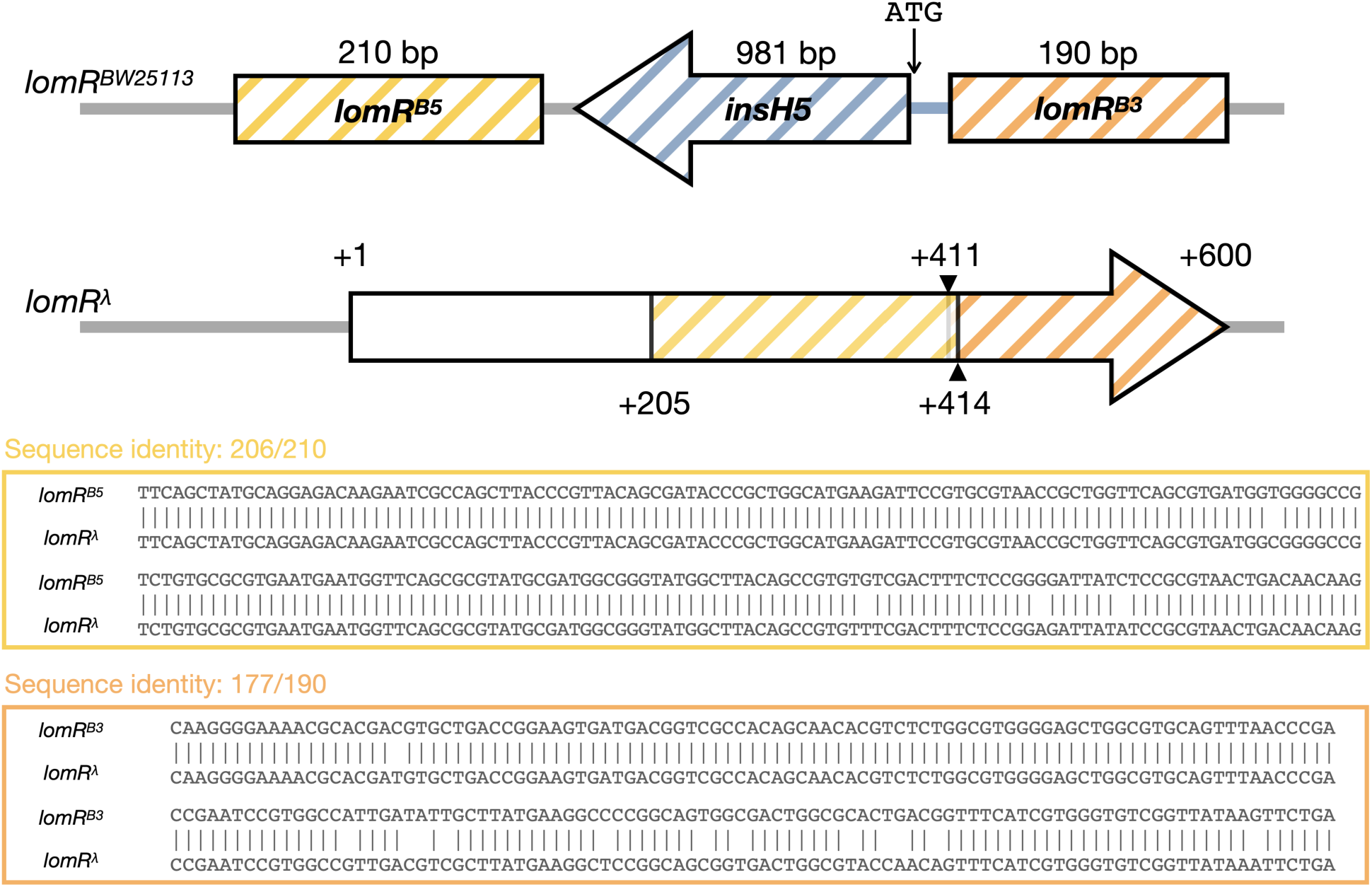
*E. coli lomR* Resembles Lambda *lomR*. (Top) *lomR^BW25113^* is similar to *lomR^λ^*. *lomR^BW25113^* is interrupted by DNA encoding an InsH5 insertion element (blue). (Bottom) DNA sequence alignment of the 5’ (yellow) and 3’ (orange) domains of *lomR^BW25113^* against those regions of *lomR^λ^*. The designations B5 and B3 denote the 5’ and 3’ coding regions, respectively, of the BW25113 *lomR* gene, as described in the main text. The DNA sequences were divided onto two lines to fit on the page. All of the lines of text read in the 5’ to 3’ direction.

**Supplementary Figure 4.**
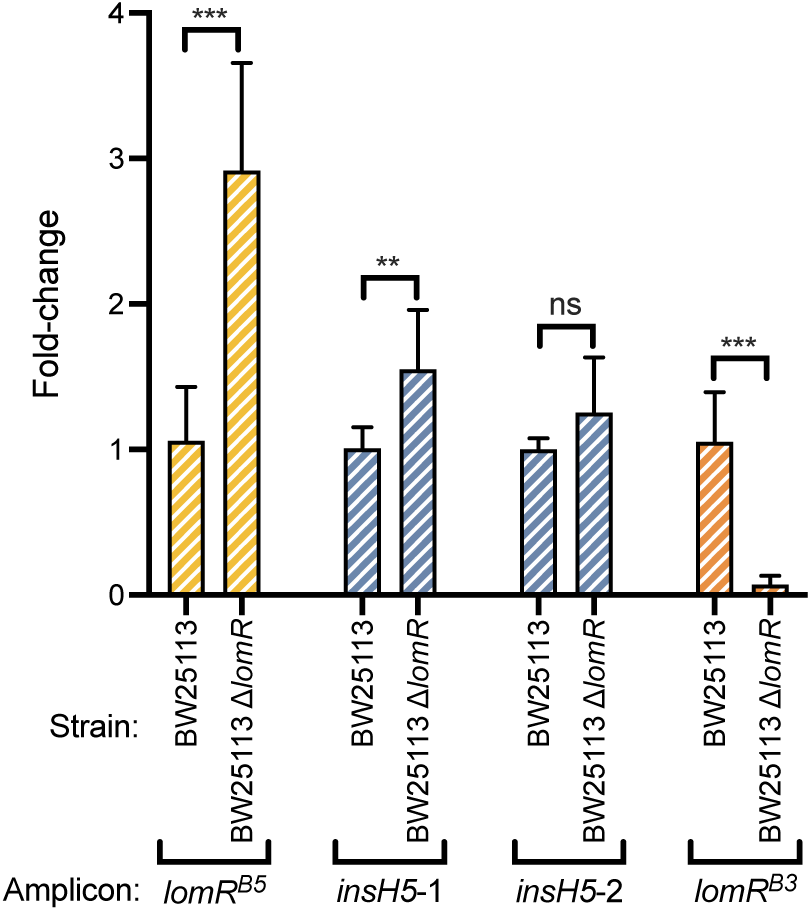
*lomR^B3^* is Not Transcribed in the BW25113 Δ*lomR* Strain. Quantitation of qRT-PCR amplification of *lomR^B5^*, *insH*5, and *lomR^B3^* in the indicated strains. *insH5*-1 and i*nsH5*-2 correspond to amplicons 2 and 3, respectively, from Figure 3C-D. Data were normalized to expression of the *gyrB* housekeeping gene. Fold-changes are relative to expression in BW25113. Data are represented as mean ± SD with *n*=3 biological and *n*=2 technical replicates. Statistical significance was calculated using one-way ANOVA Tukey’s multiple comparisons test between BW25113 and BW25113 Δ*lomR*.

**Supplementary Figure 5.**
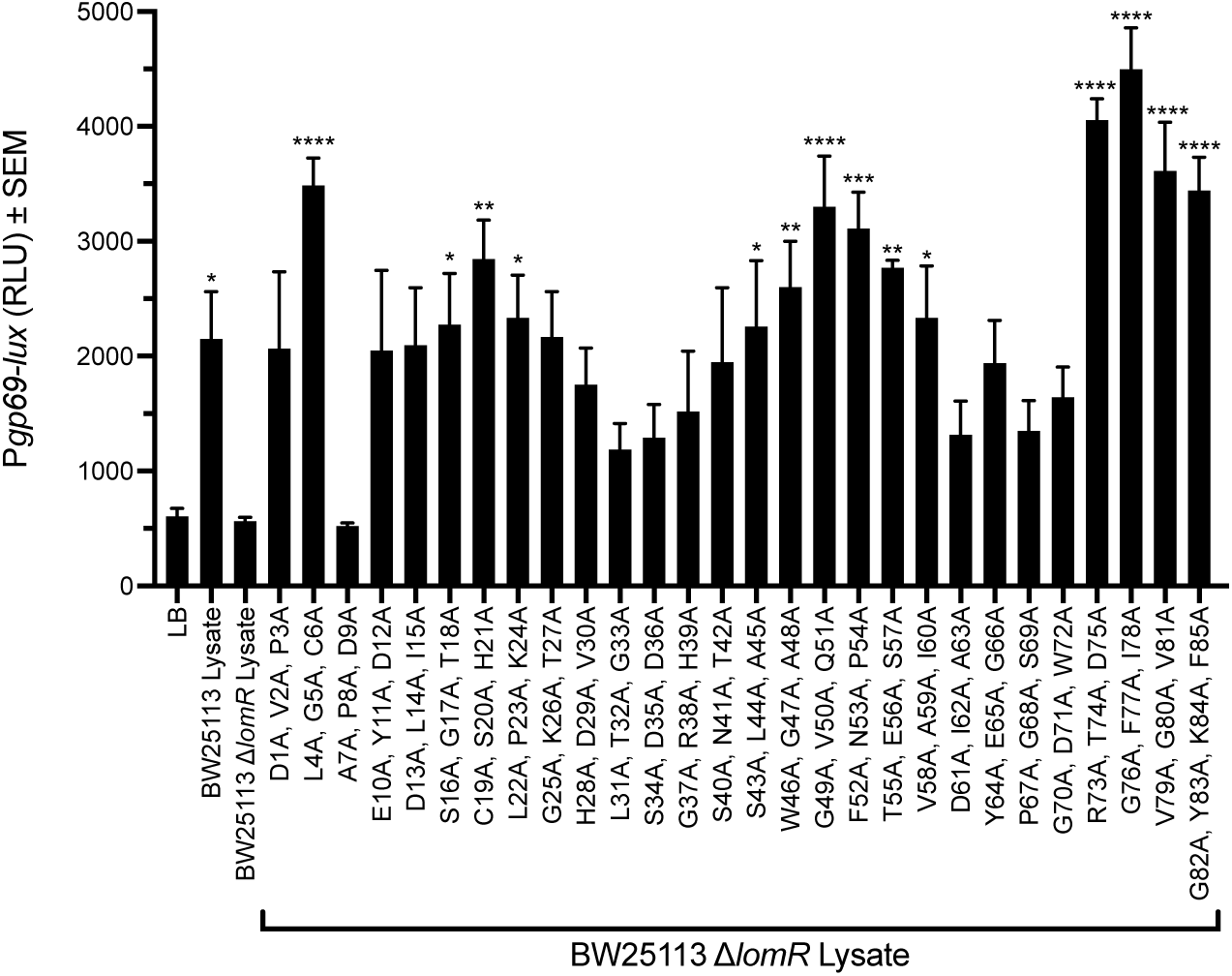
Alanine Scan of the 3’ Domain of LomR Reveals Key Residues Required for Activity. Shown are activities of LomR^B3^ in which alanine scanning mutagenesis was used to simultaneously alter three consecutive codons. This strategy involved generating 27 LomR^B3^ Ala-Ala-Ala triple mutants and 1 Ala-Ala-Ala-Ala quadruple mutant, cloning of each gene into pBAD for arabinose-inducible expression, followed by transformation into BW25113 Δ*lomR*, and preparation of lysate. All strains were grown in medium containing 0.2% arabinose prior to lysis. Assay as in Figure 2A. Data are represented as mean ± SEM with *n*=4 biological replicates. Statistical significance was calculated using one-way ANOVA Tukey’s multiple comparisons test against LB. * denotes *P*<0.05, ** denotes *P*<0.01, *** denotes *P*<0.001, and **** denotes *P*<0.0001.

**Supplementary Figure 6.**
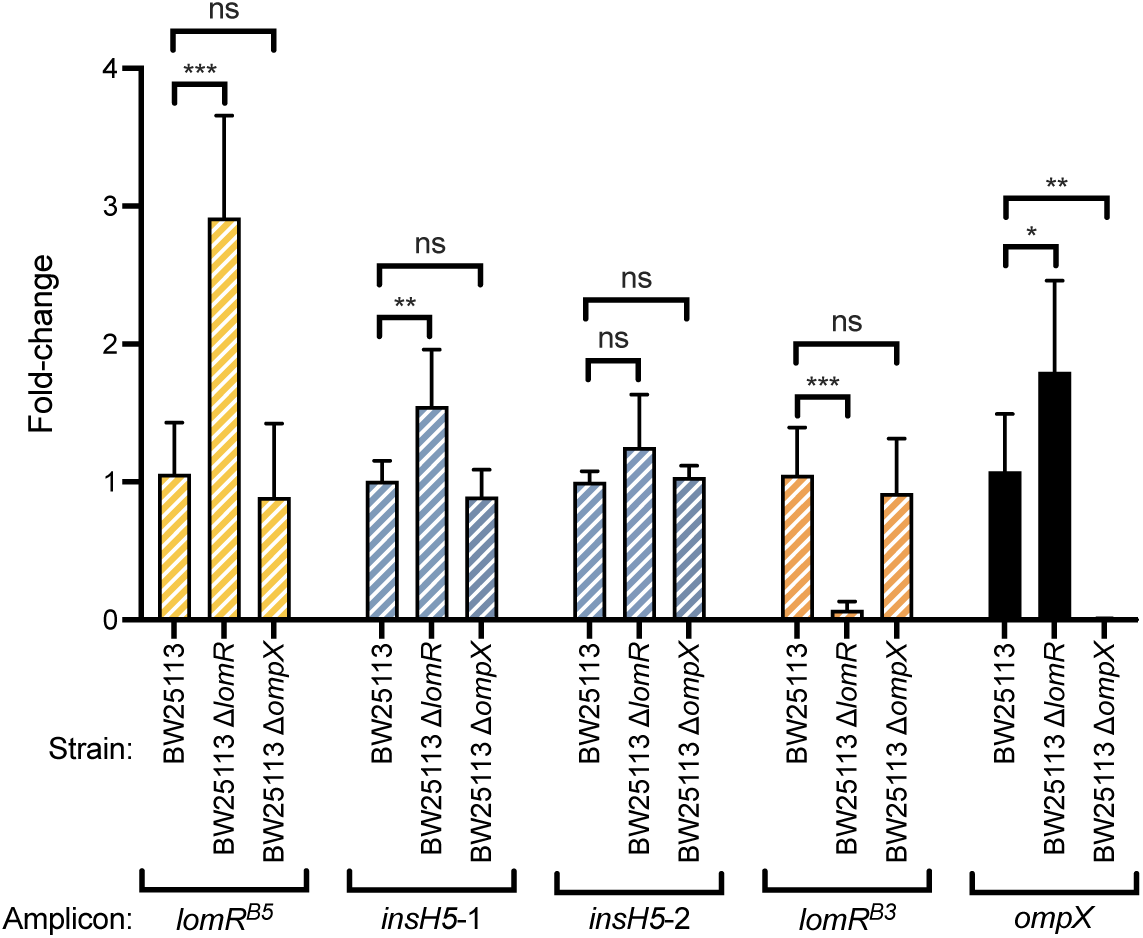
The *lomR^B3^* and *ompX^BW25113^* Genes Do Not Exhibit an Epistatic Relationship with Respect to Transcript Abundance. Quantitation of qRT-PCR amplification of *lomR^B5^*, *insH5*, *lomR^B3^,* and *ompX^BW25113^* in the indicated strains. *insH5*-1 and *insH5*-2 correspond to amplicons 2 and 3, respectively, from Figure 3C-D. Data were normalized to expression of the *gyrB* housekeeping gene. Fold-changes are relative to expression in BW25113. Data are represented as mean ± SD with *n*=3 biological and *n*=2 technical replicates. Statistical significance was calculated using one-way ANOVA Tukey’s multiple comparisons test between BW25113 and BW25113 Δ*lomR* or BW25113 and BW25113 Δ*ompX*.

**Supplementary Figure 7.**
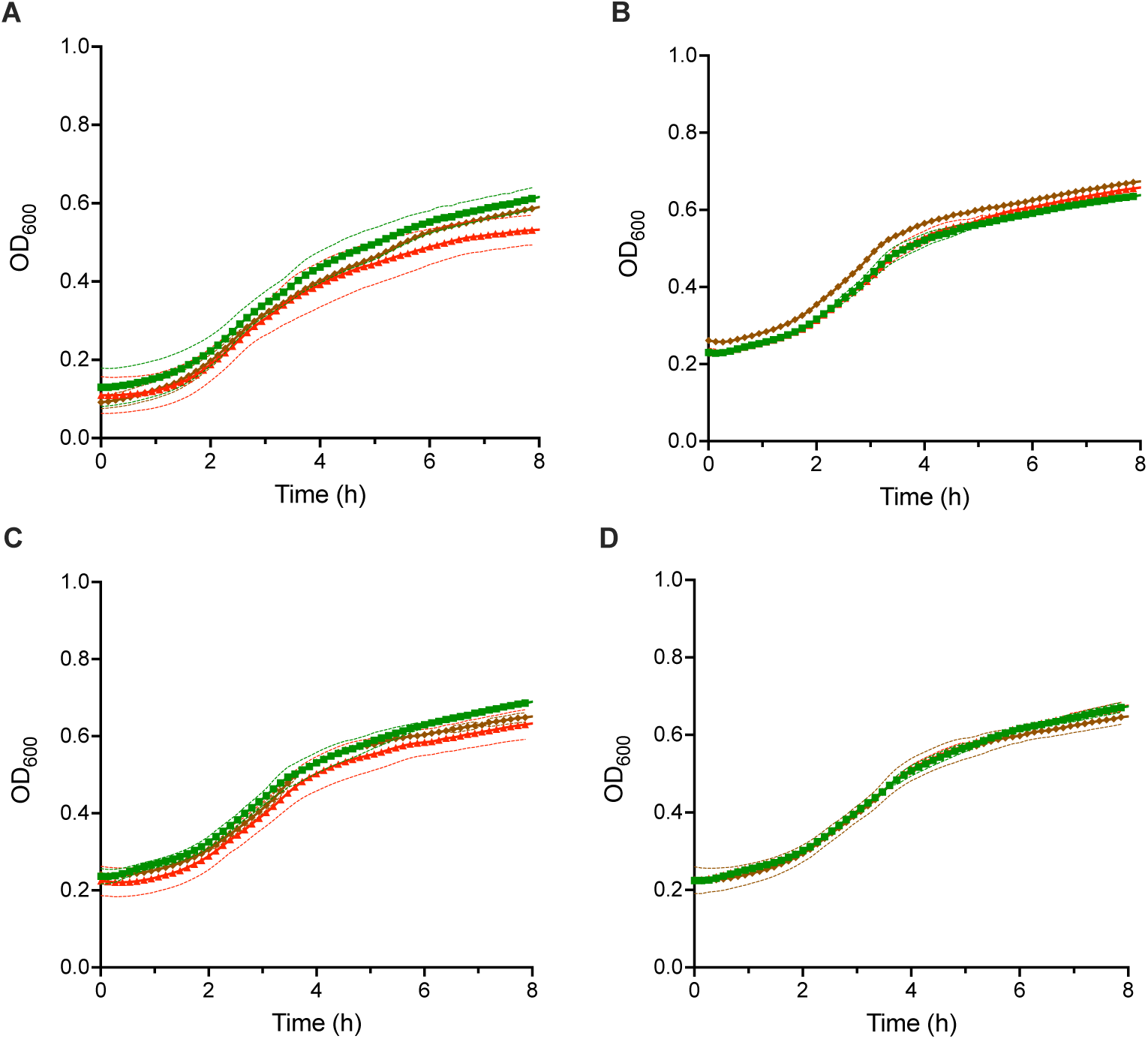
Extracellular, Not Intracellular, LomR^B3^ Induces Lysis. Growth of (**A**) BW25113 harboring control phage VP882 and arabinose-inducible *lomR^B3^*, (**B**) BW25113 harboring control phage VP882 and the pBAD plasmid, (**C**) BW25113 harboring phage VP882 *vqmA_Phage_*::Tn*5* and arabinose-inducible *lomR^B3^*, and (**D**) BW25113 harboring phage VP882 *vqmA_Phage_*::Tn*5* and the pBAD plasmid. In all growth curves, arabinose was provided at the following levels: 0.01% (green squares), 0.1% (brown diamonds), or 1% arabinose (red triangles). Data are represented as mean ± SEM (dashed lines) with *n*=4 biological replicates.

**Table S1A.**
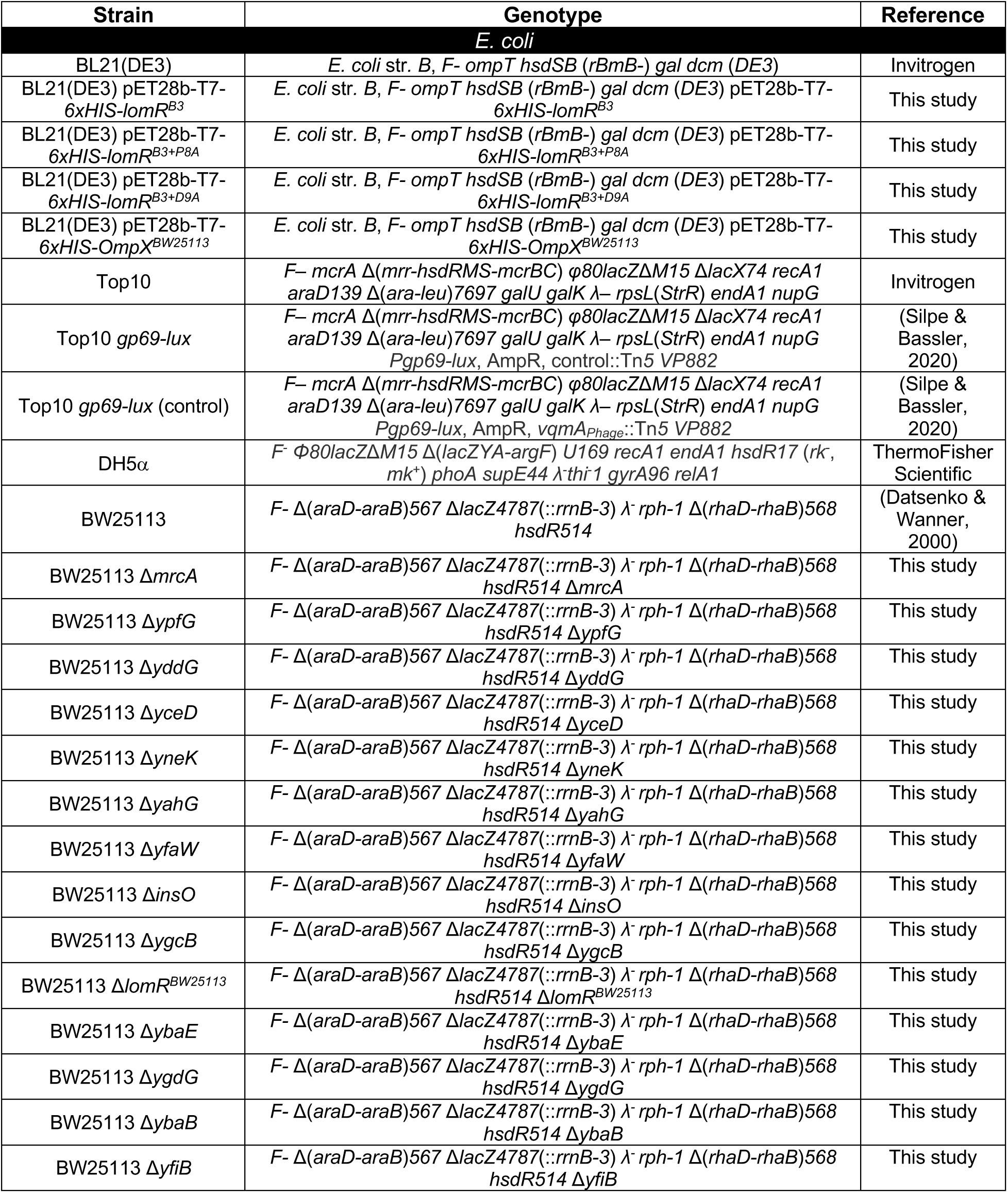
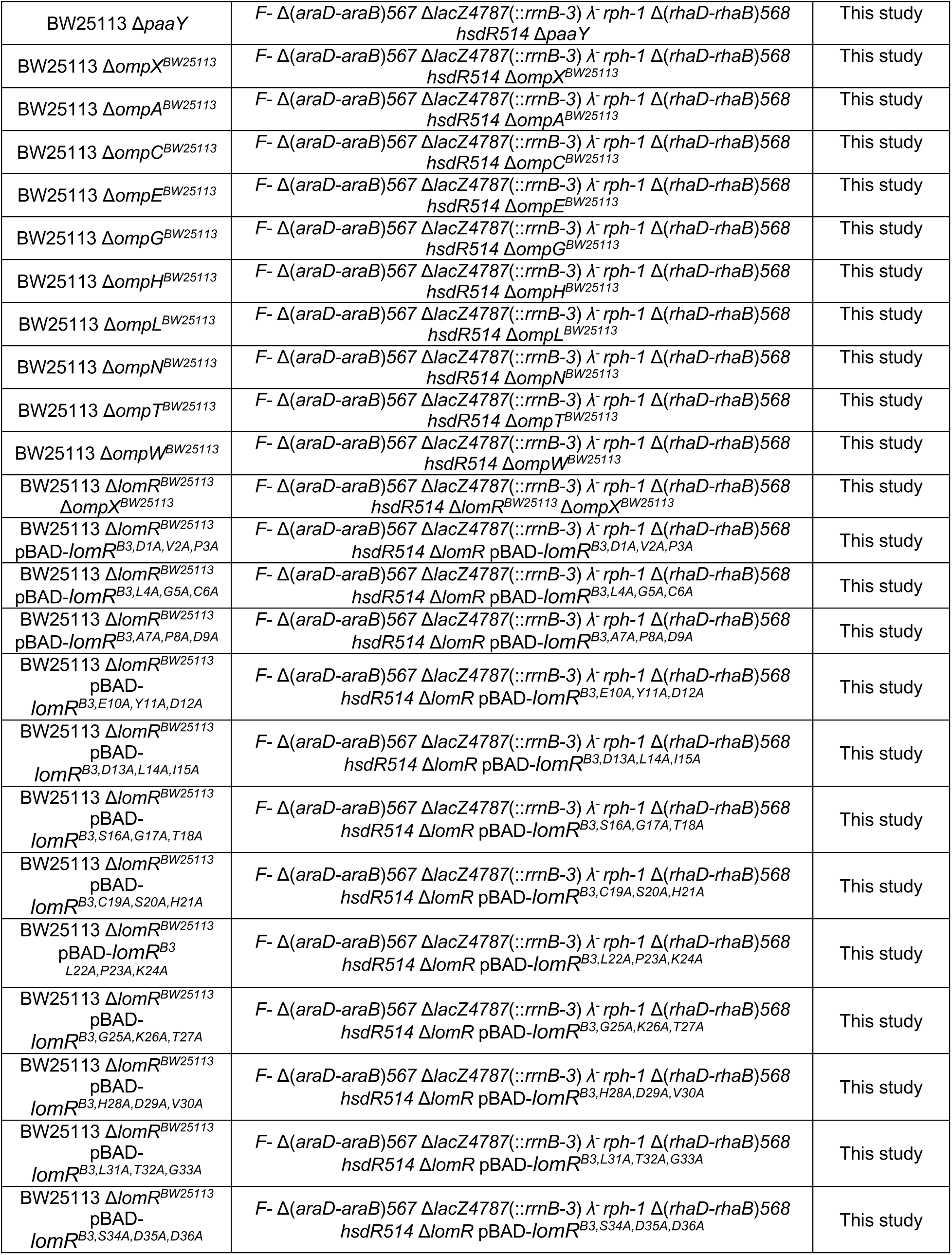
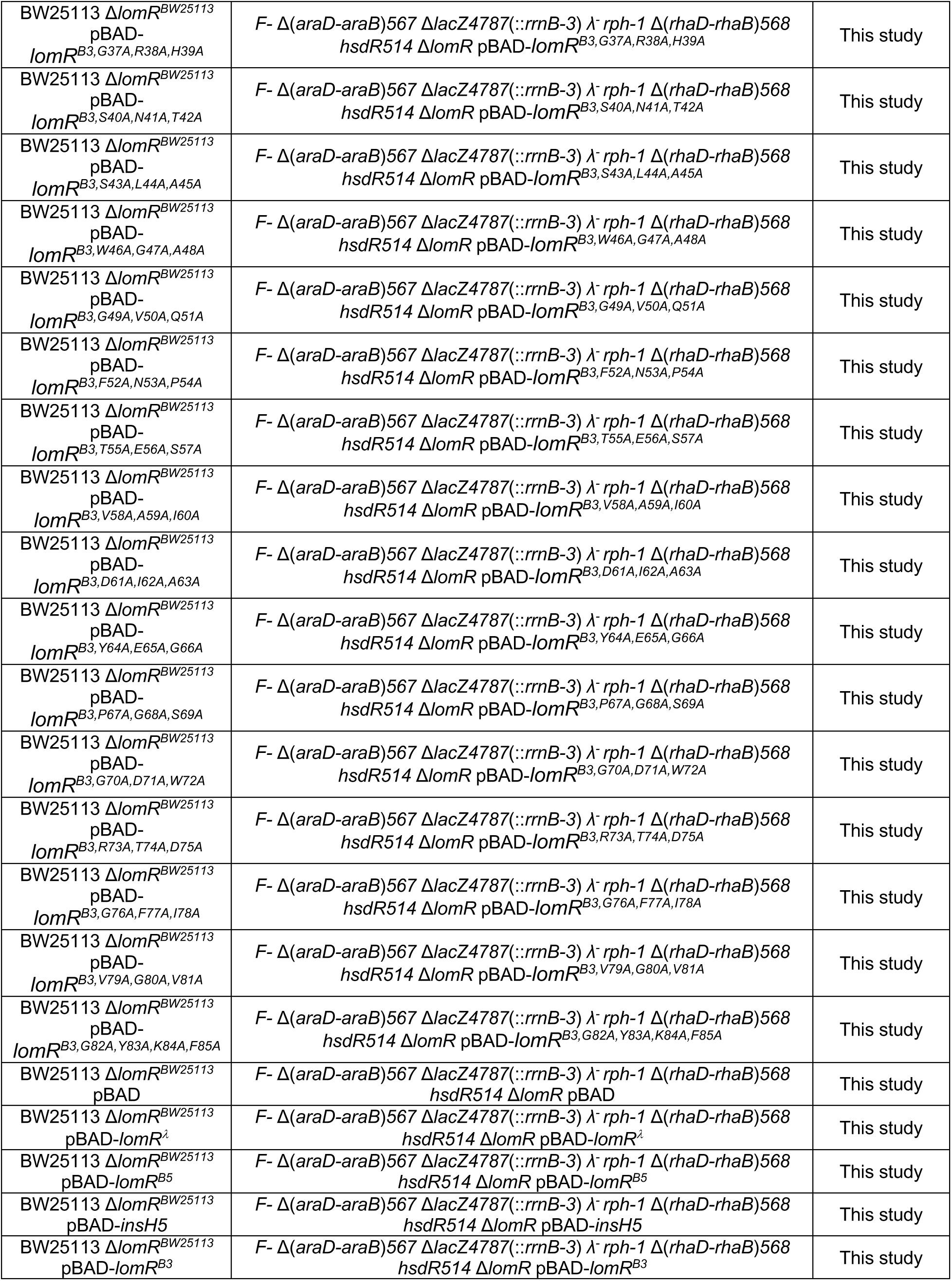
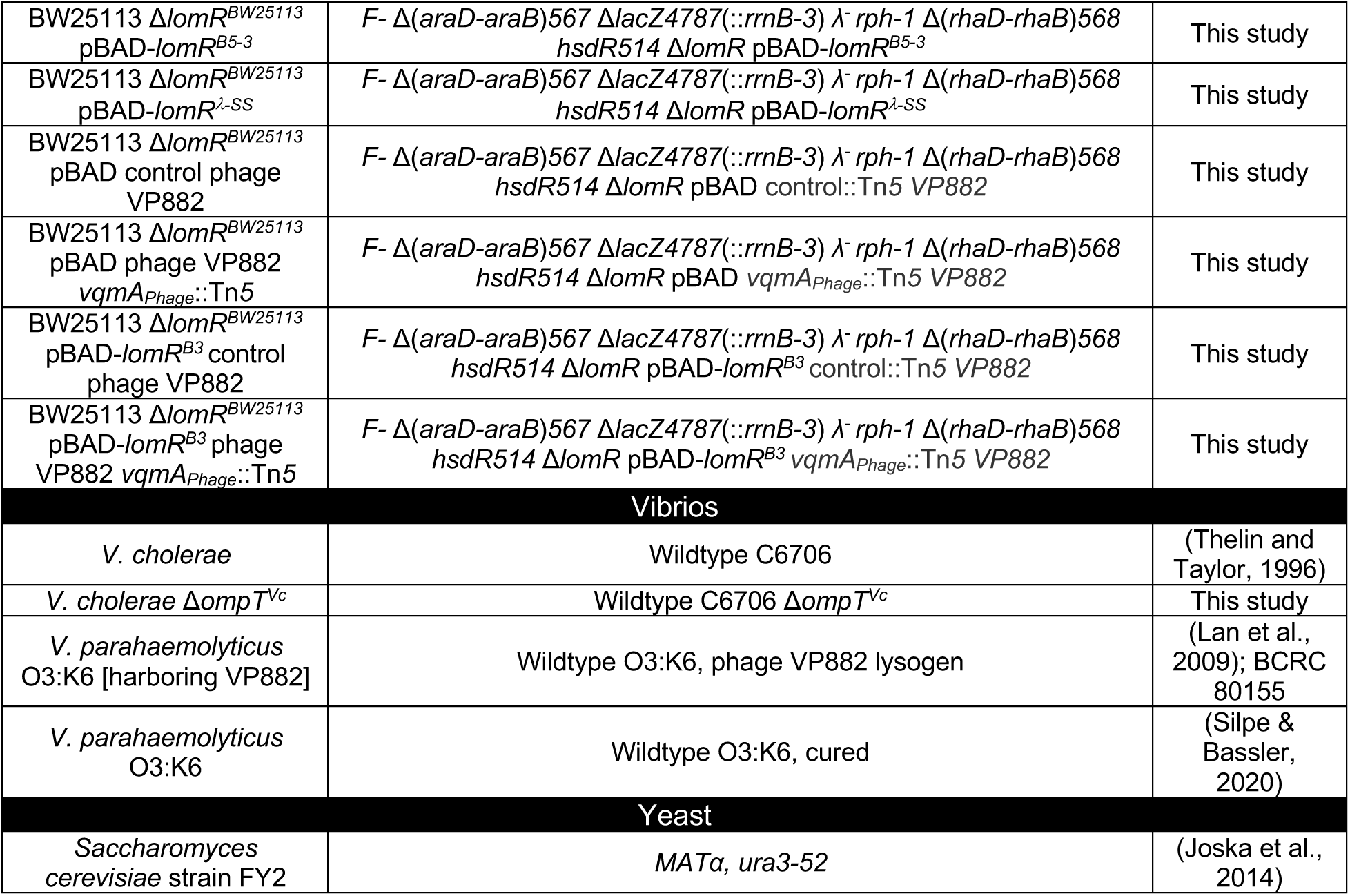
Bacterial strains used in this study.

**Table S1B.**
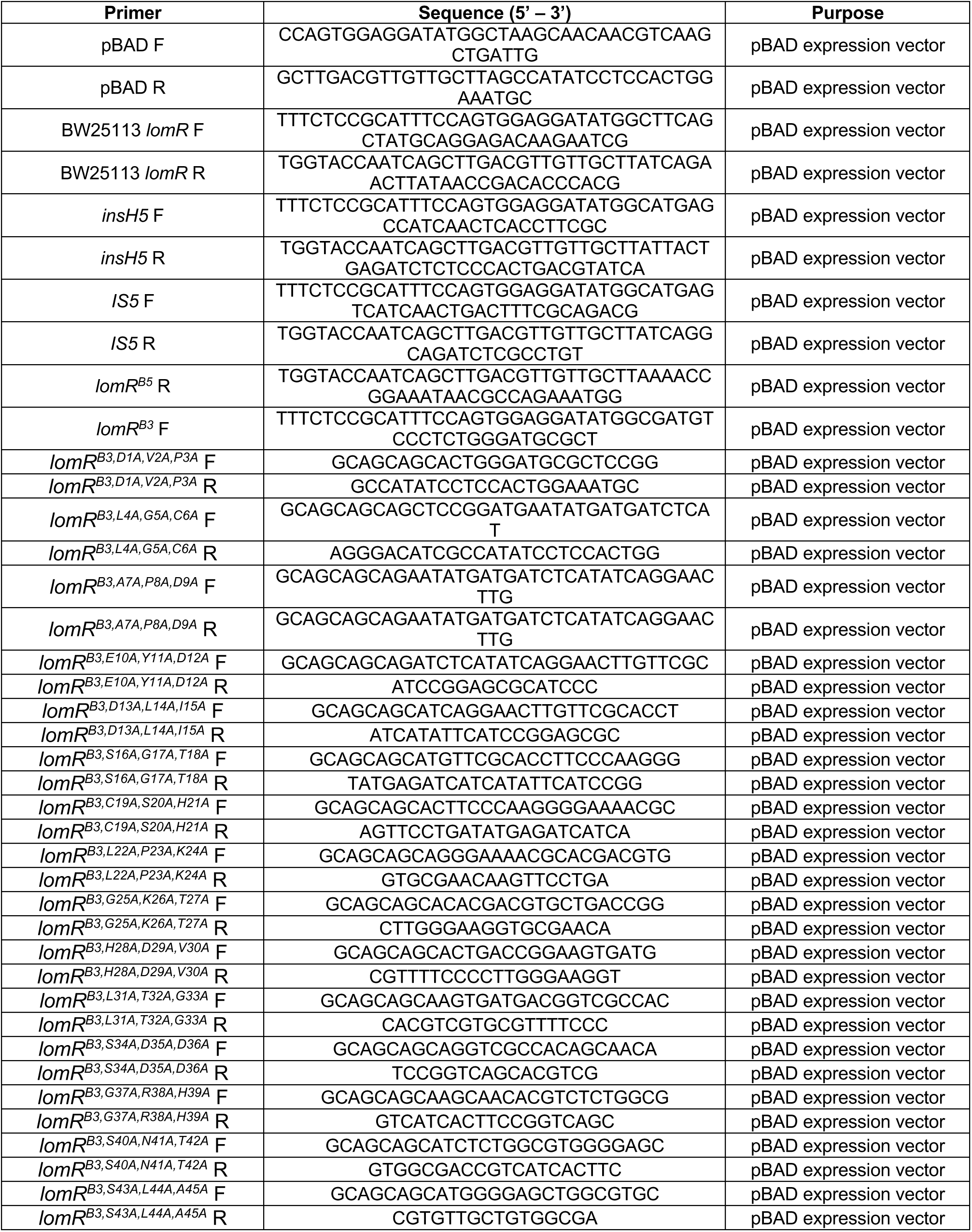
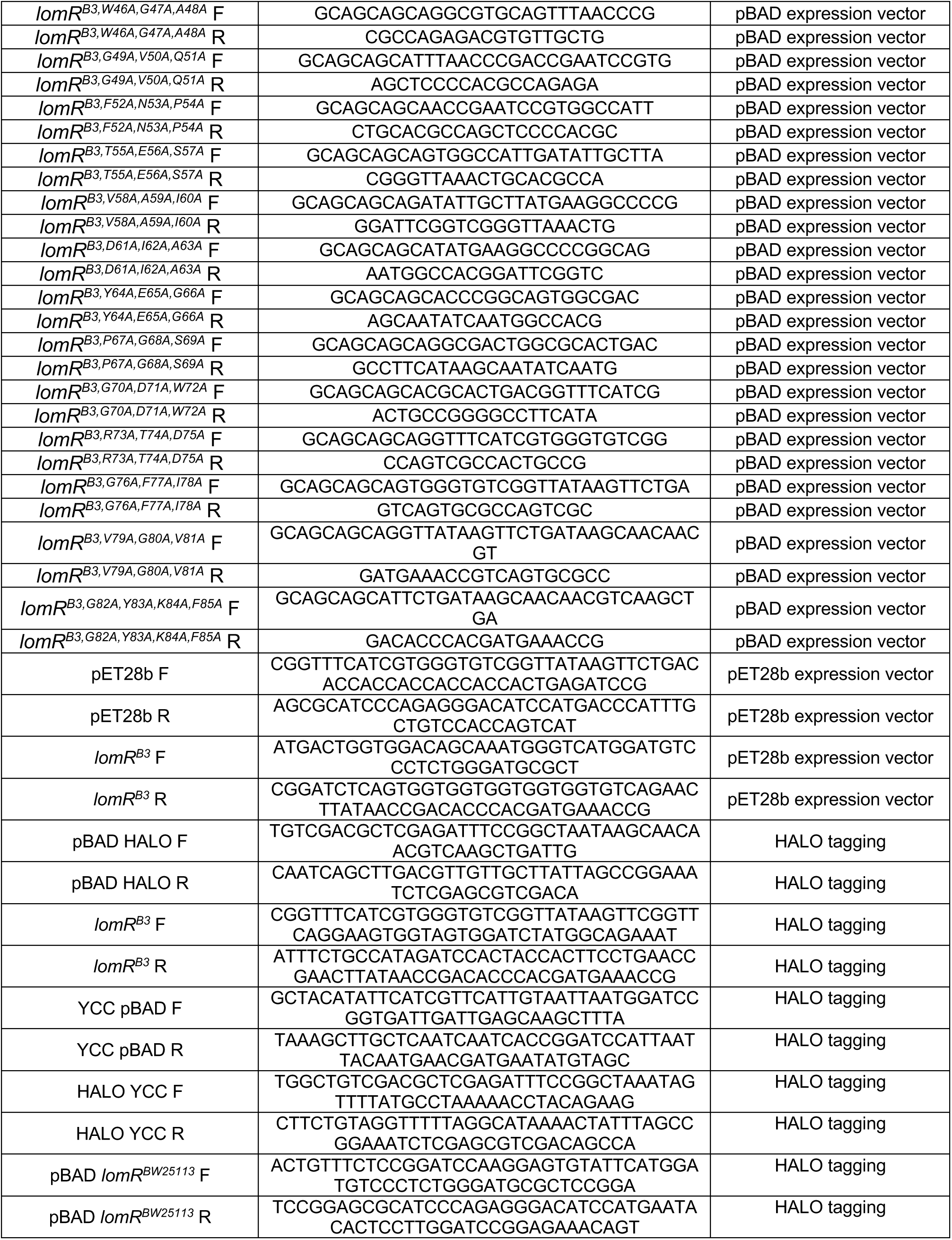
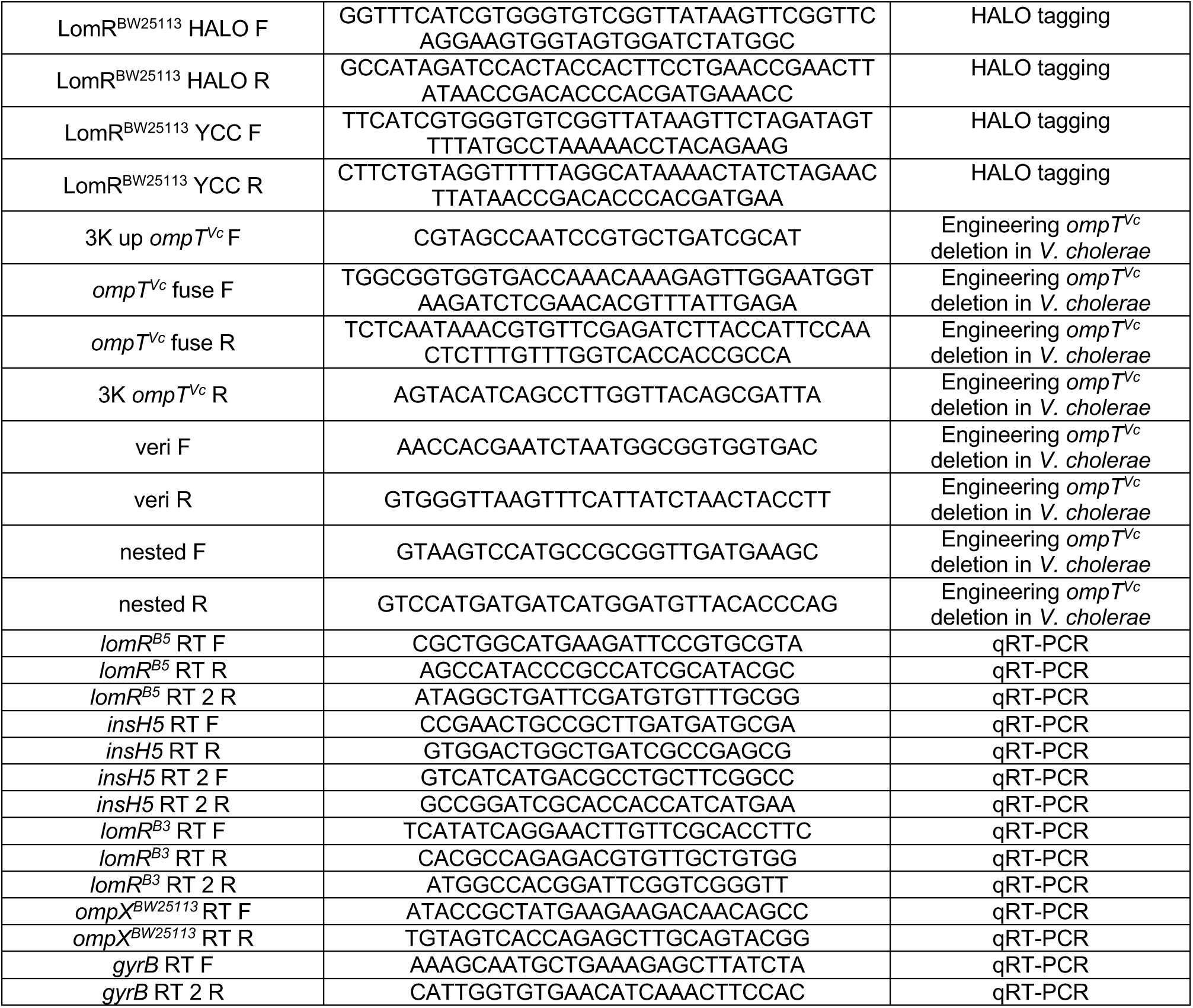
Primers used in this study.

**Table S1C.**
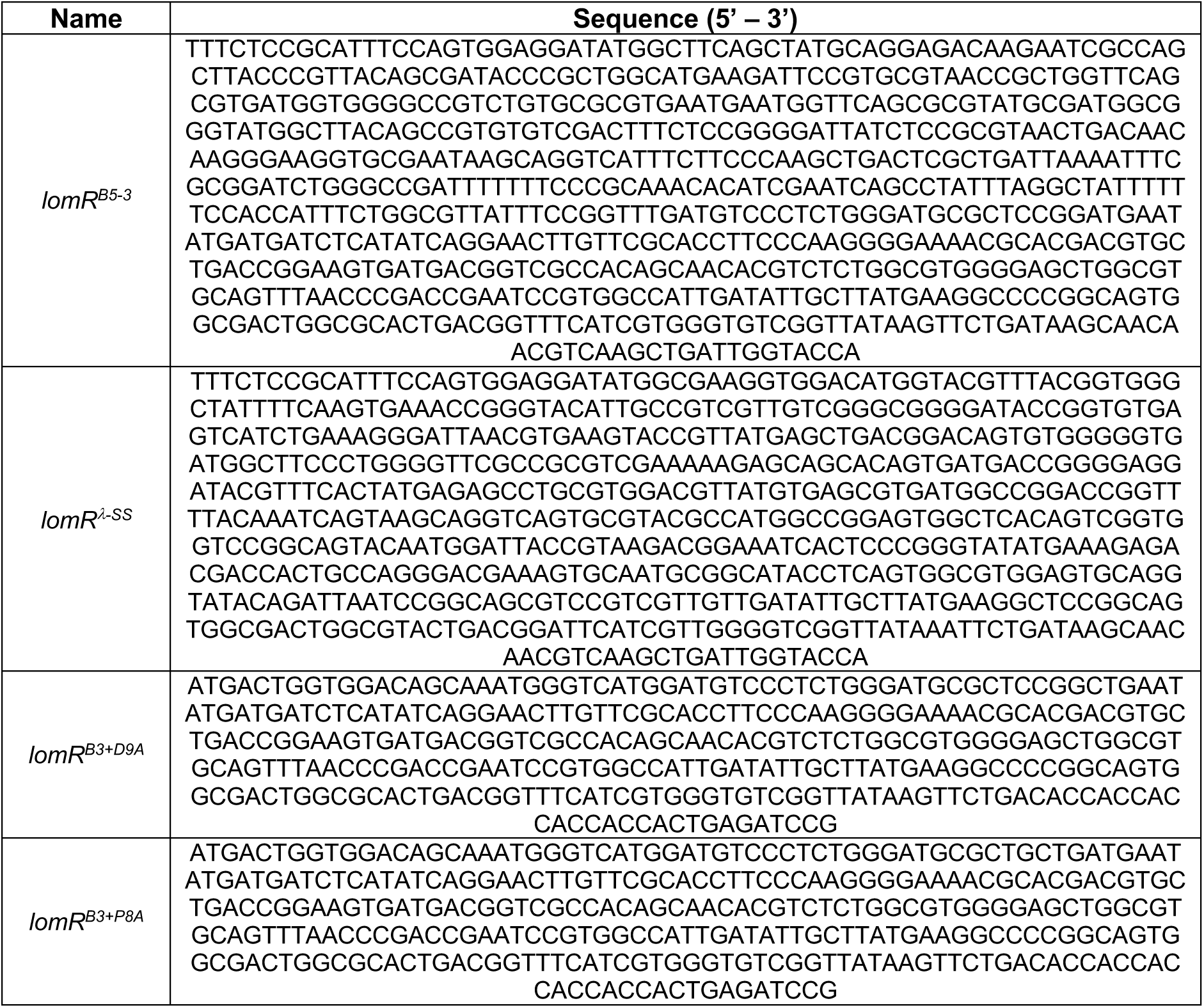
Gene blocks used in this study.

**Table S1D.**
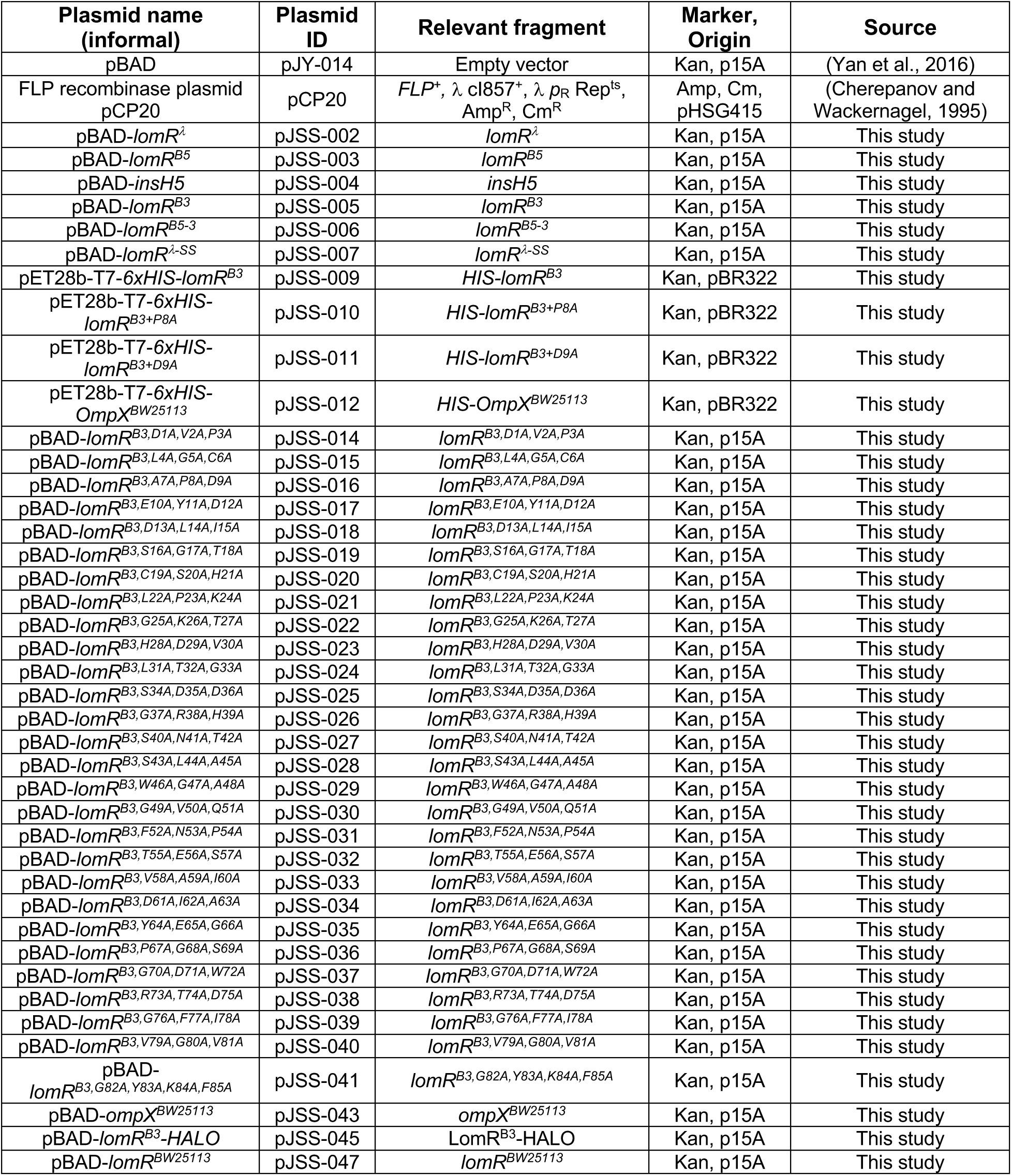
Plasmids used in this study.

